# An approximate line attractor in the hypothalamus that encodes an aggressive internal state

**DOI:** 10.1101/2022.04.19.488776

**Authors:** Aditya Nair, Tomomi Karigo, Bin Yang, Scott W Linderman, David J Anderson, Ann Kennedy

## Abstract

The hypothalamus plays a key role in regulating innate behaviors. It is widely believed to function as a system of ‘labeled lines’, containing behavior-specific neurons with characteristic transcriptomic and connectomic profiles. This view however fails to explain why, although activation of estrogen receptor-1 (Esr1) expressing neurons in the ventromedial hypothalamus (VMHvl) promotes aggression, few VMHvl neurons are tuned to attack. To address this paradox, we adopted an unsupervised dynamical systems framework to analyze population activity among VMHvl^Esr1^ neurons during aggression. We discovered that this activity contains an “integration” dimension exhibiting slow-ramping dynamics and persistent activity that correlates with escalating aggressiveness. These dynamics are implemented as an approximate line attractor in state space. Our analysis suggests a function for VMHvl to encode the intensity of behavior-relevant motive states using line attractors. This view reconciles observational and perturbational studies of VMHvl, and reveals a new mode of neural computation in the hypothalamus.

## Introduction

Innate behaviors are typically triggered by specific exteroceptive or interoceptive sensory cues that initiate a cascade of activity in the brain (Tinbergen, 1951). Cue-evoked neural activity in primary sensory or interoceptive brain regions modifies activity in downstream areas. This integration of new signals into the brain’s ongoing dynamics can drive a behavioral reaction and/or generate an internal motive state such as arousal or fear that may modify expression of future behavior. Where and how internal states are generated, and how internal states and sensory cues are jointly transformed into behavior by the brain, remains an important unsolved problem.

The hypothalamus has been implicated in the expression of innate behaviors for well over a century (Saper and Lowell, 2014). These include homeostatic behaviors such as feeding or drinking, as well as predator defense, grooming, mating, and aggression (Canteras, 2002; Hess, 1927; Hess and Brügger, 1943; Leibowitz, 1992). More recent studies have used genetically targeted optogenetic or chemogenetic perturbations to demonstrate that specific hypothalamic subpopulations control particular behaviors in a dominant manner (Anderson, 2016; Augustine et al., 2020; Chen and Hong, 2018; Hashikawa et al., 2017b; Sternson, 2013) (We use the term “dominant” to describe the effect of experimentally activating a neural population on a particular behavior, and to distinguish this population from neurons within the same brain region that may “encode” or “represent” other behavior(s) in their firing rates but which are not affected by such perturbations; see Karigo et al., 2021.) In male mice, for example, estrogen receptor type 1 (Esr1) and progesterone receptor (PR)-expressing glutamatergic neurons in the ventrolateral subdivision of ventromedial hypothalamus (VMHvl) play a dominant role in the control of aggression in male mice (Lee et al., 2014; Yang et al., 2017; Zha and Xu, 2021). Conversely, Esr1^+^ GABAergic neurons in the medial preoptic area (MPOA) play a dominant role in mating behaviors (Chen et al., 2021; Gao et al., 2019; Karigo et al., 2021; Tschida et al., 2019; Wei et al., 2018).

An important question is how the dominant behavioral influences of a given neural subclass, as identified in artificial activation experiments, relate to its population dynamics or coding properties during spontaneous occurrences of those same behaviors (Jazayeri and Afraz, 2017). In the case of MPOA, the dominant role of Esr1^+^, VGAT^+^ neurons in mating is in alignment with two other lines of investigation. First, calcium imaging experiments reveal neuronal subpopulations in the MPOA that are active during specific mating actions such as mounting or intromission (Karigo et al., 2021). And second, single cell RNA sequencing (scRNAseq) experiments have identified transcriptomically distinct MPOA neuronal subpopulations that express c-fos during specific reproductive social behaviors such as mating (Moffitt et al., 2018) (Figure 1A). Thus, in the case of MPOA, both observational and perturbational studies align to support a key role for Esr1^+^ GABAergic neurons in the control of specific mating-related actions.

**Figure 1:**
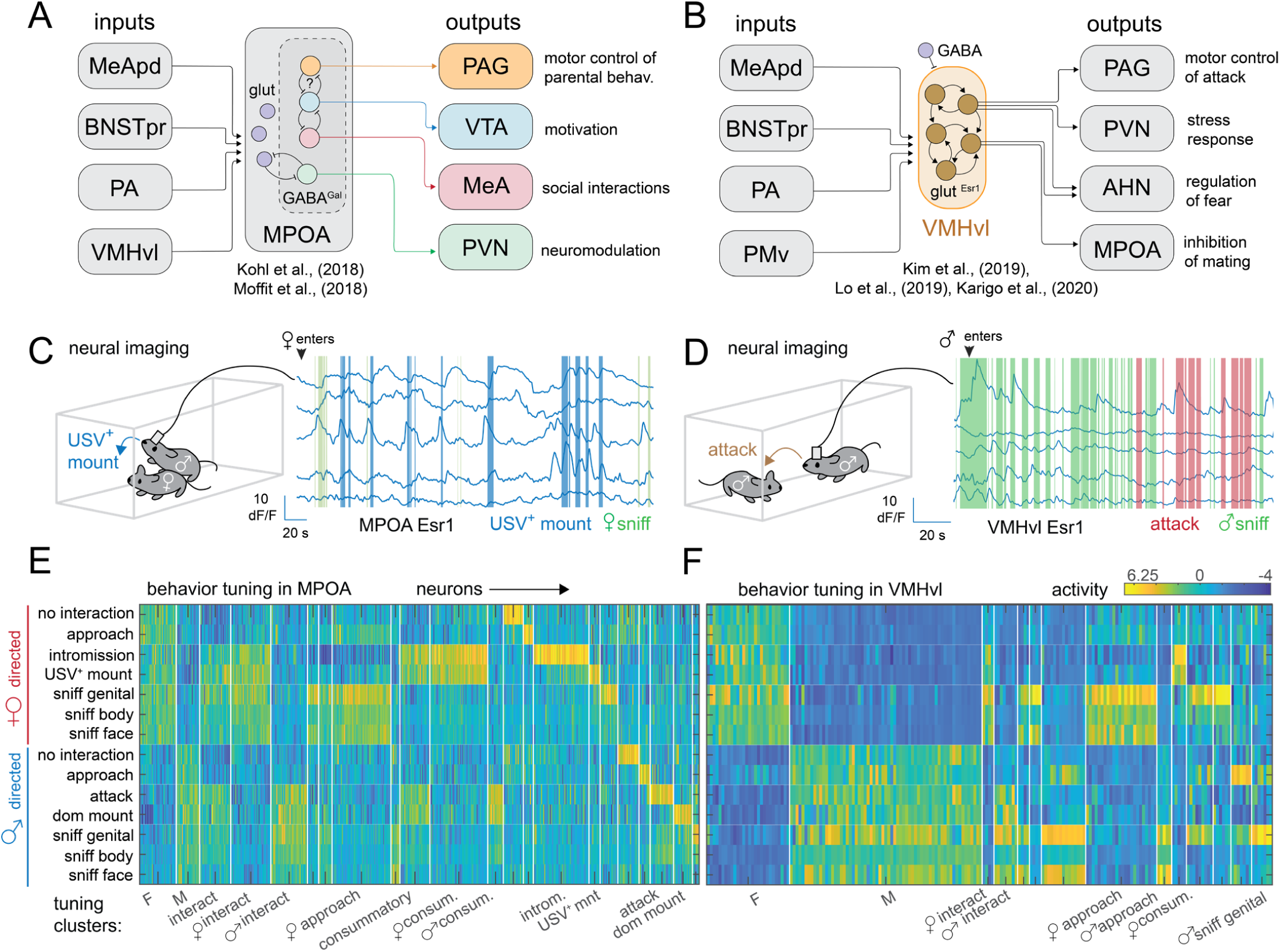
Cytoarchitectures and cellular representations in a neural system regulating social behavior. A: cytoarchitecture of the MPOA. B: cytoarchitecture of the VMHvl. C: neural imaging studies of Esr1+ neurons in the MPOA find neurons tuned to mounting behavior. D: neural imaging studies of Esr1+ in the VMHvl find few if any neurons tuned to aggressive behavior. E: clustering of recorded Esr1+ neurons in MPOA using a regression model reveals many populations of behavior-tuned neurons (n =306 neurons from 3 mice). F: clustering of recorded Esr1+ neurons in VMHvl using a regression model reveals that most neurons are tuned to the sex of the intruder and few neurons are tuned to any behavior (n = 391 neurons from 4 mice).

In contrast, while manipulations of VMHvl^Esr1/PR^ neurons have revealed a dominant role for these neurons in aggression, there is only a weak correspondence between VMHvl^Esr1^ neuron activity and specific occurrences of aggressive behavior; instead, most individual VMHvl^Esr1^ neurons exhibit strong selectivity for either male or female conspecifics (Remedios et al., 2017). Although some neurons show weak selective modulation during aggression, mating, or investigative behavior, most cells exhibit mixed selectivity for multiple behaviors (Karigo et al., 2021; Remedios et al., 2017). Furthermore, although VMHvl^Esr1^ neurons exert a dominant influence on aggression, they are also required (albeit not sufficient) for normal levels of mating (Karigo et al., 2021; Yang et al., 2013). These data suggest that the function of the male-vs. female-selective VMHvl^Esr1^ neurons, as revealed by observational experiments, may be to encode conspecific sex identity (Yang and Shah, 2014), or an internal motive state that is strongly correlated with conspecific sex, i.e., aggressiveness (Falkner et al., 2014; Falkner et al., 2016) vs. mating drive (Zhang et al., 2021). Because male mice typically exhibit aggression towards males and mating towards females, intruder sex and behavior are highly correlated, making it difficult to distinguish these alternatives. (However a subset of VMHvl^Esr1^ neurons in female mice has been shown to promote aggression towards both males and females; (Hashikawa et al., 2017a; Liu et al., 2022)).

The failure to identify a robust encoding of aggressive behavior at the level of single neurons in VMHvl seems paradoxical given the high specificity of optogenetic and chemogenetic activation for aggressive behaviors, including dominance mounting and attack (Karigo et al., 2021; Lee et al., 2014; Yang et al., 2017). It is especially puzzling in light of the strong correspondence between optogenetic phenotypes and behavior-specific single-cell tuning observed in MPOA ((Karigo et al., 2021; Kohl et al., 2018; Moffitt et al., 2018; Wu et al., 2014) This paradox has led us to investigate whether population-based, rather than single-neuron, coding might underlie the control of aggression by VMHvl^Esr1^ neurons.

Here we use recurrent switching linear dynamical systems (rSLDS) (Linderman et al., 2017) analysis to investigate whether the dynamics of VMHvl^Esr1^ population activity can reveal an encoding of aggressive behavior and/or drive in this nucleus. This analysis uncovered an approximate line attractor in neural activity space, progression along which is correlated with the intensity and type of aggressive behavior displayed. By contrast, in MPOA we observe rotational rather than line attractor dynamics, consistent with our observation of the sequential activation of behaviorally tuned MPOA neurons during mating. Taken together, our results reveal that different hypothalamic nuclei use different neural coding schemes to control closely related social behaviors. They also provide evidence for line attractors as a mechanism by which subcortical circuits may generate persistent and scalable internal states that shape the expression of survival behaviors.

## Results

### Cellular tuning analysis reveals behaviorally selective neural populations in MPOA but not in VMHvl

Although optogenetic stimulation of VMHvl^Esr1^ glutamatergic or MPOA^Esr1^ GABAergic neurons evokes aggression and mating, respectively (Hashikawa et al., 2017a; Karigo et al., 2021; Lee et al., 2014; Wei et al., 2018; Yang et al., 2013), calcium imaging during social interaction reveals more complex patterns of neuronal activation (Figure 1A, B) (Karigo et al., 2021; Remedios et al., 2017). To illustrate these differences more clearly, we compared behavioral tuning at the level of individual Esr1^+^ neurons in VMHvl vs. MPOA during dyadic social interaction. Specifically, we re-analyzed calcium imaging data (Karigo et al., 2021) recorded from Esr1^+^ neurons in these nuclei using a head-mounted microendoscope (Ghosh et al., 2011; Ziv et al., 2013), in sexually experienced male C57Bl/6N*^Esr1-2A-Cre^* mice during standard resident-intruder assays, using male or female BalbC intruders (Figure 1C, D).

To compare the responses of Esr1^+^ neurons in VMHvl and MPOA, we computed the mean activation of each imaged neuron during 14 different, manually annotated behavioral actions and grouped neurons by their normalized pattern of activation across all actions (VMHvl: N= 306 neurons from 3 mice; MPOA: N= 391 neurons from 4 mice.) This analysis revealed that many MPOA clusters contained neurons only active during specific social behaviors, such as intromission, ultrasonic vocalization-accompanied (USV^+^; reproductive) mounting, attack, and USV^-^ (dominance) mounting (Figure 1E). A relatively small proportion was activated in a sex-specific but not behaviorally specific manner. In contrast, most neurons in VMHvl were broadly activated in an intruder sex-specific manner, with very few neurons showing preferential activation during specific behavioral actions (Karigo et al., 2021; Remedios et al., 2017) (Figure 1F). Most strikingly, we observed a higher proportion of attack-specific neurons in MPOA than in VMHvl, despite the former’s dominant influence on mating.

This analysis indicates that MPOA and VMHvl cannot simply be considered analogous nuclei that govern opponent social behaviors in comparable fashion. MPOA contains multiple distinct behaviorally-tuned subpopulations, consistent with c-fos - analysis that shows that this nucleus contains specific transcriptomic cell-types activated during different social behaviors (Kohl et al., 2018; Moffitt et al., 2018). In contrast, VMHvl contains very few neurons tuned to specific behaviors, consistent with the paucity of c-fos-labeled action-specific transcriptomic cell types in VMHvl (Kim et al., 2019). This is also consistent with the conclusions from our earlier imaging studies of VMHvl^Esr1^ neurons, which examined a different dataset using choice probability analysis (Remedios et al., 2017).

### Unsupervised dynamical systems analysis of neural activity during social behavior

Our analysis of single cell tuning among VMHvl^Esr1^ neurons presented a paradox: optogenetic stimulation of these cells elicits specific social behaviors (aggressive sniffing, USV^-^ mounting and attack (Karigo et al., 2021; Lee et al., 2014)), yet we observed only a few cells (<10%) that are selectively active during these specific behaviors ((Karigo et al., 2021; Remedios et al., 2017) and this study). However, the lack of neurons exclusively activated during a particular behavioral action such as attack does not mean that VMHvl contains no information about ongoing behavior. Indeed, we previously demonstrated that the occurrence of social behavior actions, including attack and investigation, could be decoded from the activity of VMHvl neurons (Remedios et al., 2017). Nevertheless, that analysis did not provide insight into how VMHvl population dynamics control behaviors or shape the transitions between them.

In other neural systems, population analysis via fit dynamical systems has helped identify the neural encoding of values that are not apparent in neuron-by-neuron analysis (Hulse and Jayaraman, 2020; Shenoy et al., 2013; Vyas et al., 2020). We therefore investigated whether behavioral representations among VMHvl^Esr1^ neurons might be encoded at a population level, using an unsupervised dynamical systems approach.

To do so, we fit a dynamical model to the population activity of VMHvl^Esr1^ cells from each of several (n=6) individual mice over the course of an entire male-male or male-female encounter (duration 5.1± 0.68 minutes for male-male encounters and 11.4± 0.68 minutes for male-female encounters) Specifically, we fit a recurrent switching linear dynamical system (rSLDS) (Linderman et al., 2017), a class of model that approximates a complex non-linear dynamical system using a composite of linear dynamical systems. The rSLDS model offers three features that make it useful for further analysis. First, by “switching” between linear systems, the model can approximate the dynamics of a complex nonlinear system, while still permitting the fit model to be analyzed using the abundance of techniques available for linear systems (Linderman et al., 2019). Second, rSLDS can model the contributions of both external input and intrinsic dynamics to the evolution of neural population activity. The intrinsic component of the fit rSLDS model describes how neural population activity is expected to evolve over time in the absence of external perturbation; this can be visualized as a “flow field” of the system. And third, the “state” of the rSLDS, defined as the linear system that best fits the neural population activity and dynamics at a given time, can be used to segment neural activity into distinct regimes in an unsupervised manner, without incorporating any behavioral data. States identified by rSLDS can subsequently be aligned to behavioral data recorded concurrently with neural population activity, to determine whether neural states correspond to specific behavioral bouts or epochs (Supplemental Figure S1A).

rSLDS operates under the assumption that a substantial fraction of the observed variance of recorded neural activity can be captured by a low-dimensional set of latent variables, which it infers from that activity. We initialized the latent variables using Factor Analysis, selecting the number of factors for each mouse as the fewest needed to capture ≥90% of observed variance. The model segments population activity in this low-dimensional space into a set of discrete states, where the likelihood of the neural population being in a given state is determined solely by its position in this space (Figure 2A, step ①-②). Linear dynamical systems are fit to the latent variable dynamics within each discrete state, and model parameters are refined using variational inference to obtain the final model (Figure 2A, step ③). Each state, defined by its own dynamics matrix, dictates how the latent variables underlying neural activity evolve in time from any given point in that particular state space, as well as how that evolution is modified by external input. Finally, to visualize the fit system, we computed its first two principal components, and plotted the inferred dynamics in that space either as a flow field, wherein arrows show the direction and rate of change of activity at each point, or as a 3D landscape, where the rate of change in activity is converted to the height of a 3D landscape (Figure 2A step ⩽ right).

**Figure 2:**
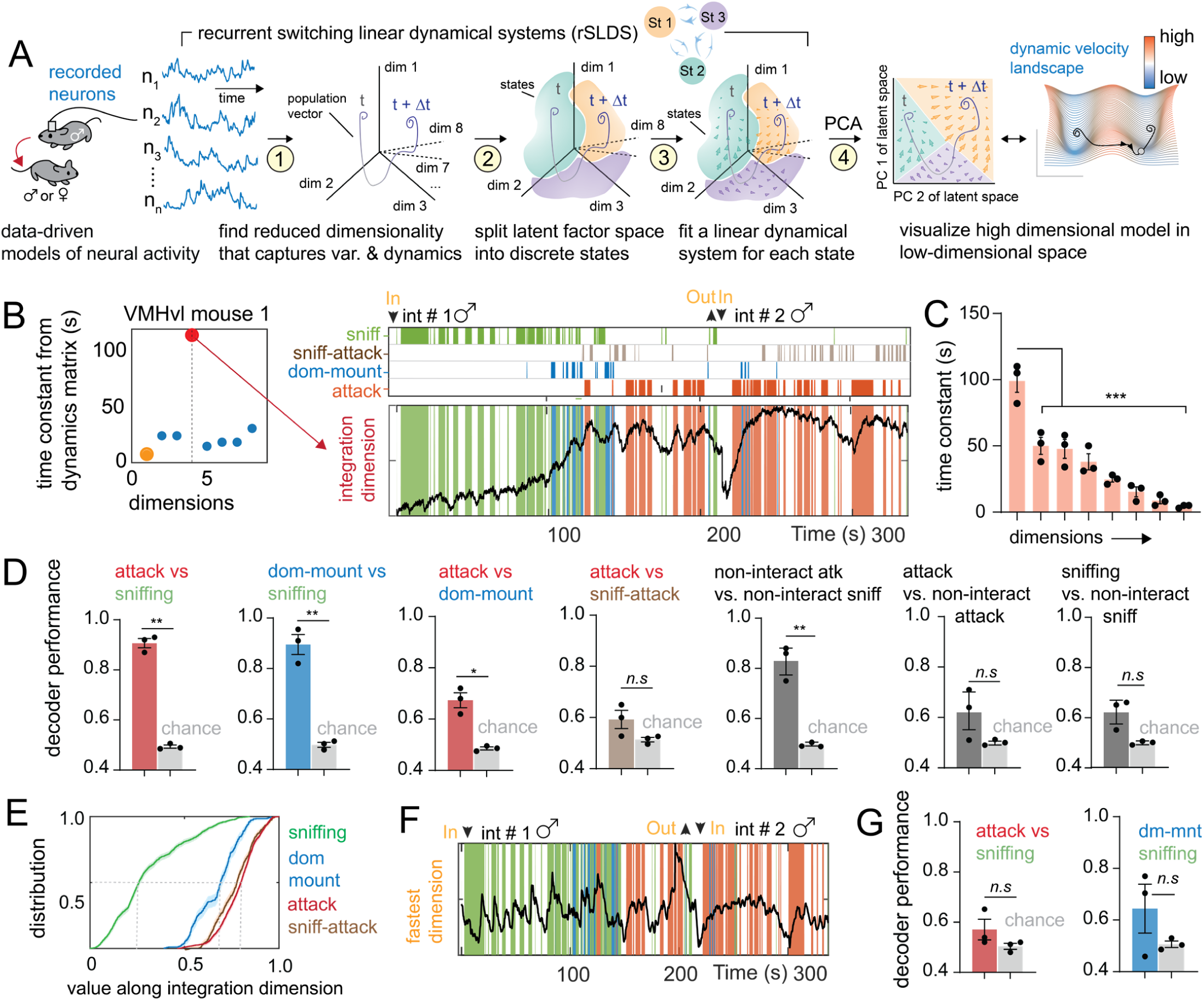
Dynamical analysis of VMHvl neural activity reveals an integrator dimension that correlates with aggressive escalation. A: application of recurrent switching linear dynamical systems (rSLDS) to neural data obtained from the MPOA or VMHvl. B: time constants of dynamics matrix of attack enriched state from example VMHvl mouse 1. The red dot highlights the largest time constant or integration dimension. C: Time constant from all animals arranged in decreasing order. The integration dimension (dim 1) is significantly larger than all other dimension (p < 0.001, n = 3 mice). D: decoding behaviors from integration dimension (**p < 0.005, *p<0.01, n = 3 mice). E: empirical cumulative distribution of value of integration dimension (normalized) for various behaviors. F: temporal dynamics of fastest dimension in example VMHvl mouse 1. G: decoding behaviors from fastest dimension across animals (n = 3 mice).

We found that 7-8 latent dimensions could capture 90% (7.3± 0.3, N=4 mice) of observed variance in VMHvl, suggesting that a relatively low-dimensional latent variable model provided a reasonable approximation of neural activity in this region. In fitting our model, we set the number of states to be used and the dimensionality of the neural system that maximized the likelihood of the data, determined using cross validation in each mouse separately (Supplemental Figure S1B-E). As input to the model, we used the distance between animals and their facing angle (orientation of the resident towards the intruder) as a proxy for the strength of social sensory cues (Falkner et al., 2014; Segalin et al., 2021). This 2D sensory signal is fed as input to the latent variables, scaled linearly by a fit set of weights to allow each latent variable to be driven to different extents by external input. Biologically, input to a given latent variable can be interpreted as that input activating (or inhibiting) the neurons that contribute strongly to that variable.

### Dynamical systems analysis of VMHvl neural activity reveals an integrator dimension that correlates with aggressive escalation

We first examined the dynamics of rSLDS models fit to VMHvl^Esr1^ neurons during encounters with male intruders. Our best fit models required either three or four rSLDS states; interestingly, attack behavior mostly occurred during a single rSLDS state in all animals (state 3, Supplementary Figure S1F-H). Attack behavior was not tightly linked with the onset of this state, but rather the probability of exhibiting attack was elevated within the state, and on average peaked 20 seconds after the state’s onset across animals (state-triggered average, N = 3 mice, Supplemental Figure S1F_4_). Epochs of the state enriched for attack also lasted much longer than individual bouts of attack within each epoch (state 3 epoch duration: 83.5± 7.5 seconds, attack bout duration: 3.66± 0.44 seconds, N = 3 mice, Supplemental Figure S1F_5_, G_3_. H_3_). In order to better understand neural population dynamics related to attack behavior, we examined the dynamics matrix for this state, which describes how neural activity in the latent variable space changes with time.

The eigenvalues of the dynamics matrix reflect the rate at which orthogonal modes (dimensions) of activity in the system decay to zero following external input, and are referred to as the time constants of these dimensions (Maheswaranathan et al., 2019; Strogatz, 2018). External input received along dimensions with short time constants will quickly decay to zero, whereas input to dimensions with long time constants persists and decays slowly. Examining the time constants of the dynamics matrix of the attack-related state revealed a single dimension with an estimated time constant of over 100 seconds (Figure 2B, Left, red dot). Because systems with long time constants approximately integrate their input over time, we refer to the longest time constant dimension the “integration” dimension. Activity along the integration dimension ramped up at the onset of dominance mounting (a low-intensity aggressive behavior, (Karigo et al., 2021)), and persisted as the animal attacked (Figure 2B, Right). A single integration dimension, with a time constant significantly higher than that of the other latent dimensions, could be found in the rSLDS model for each animal (N=3 mice, Figure 2C). The integration dimension accounted for 17.24%± 1.9% of overall variance of neural activity across animals (N = 3 mice). This is significantly higher than the variance explained by an “attack-decoding” dimension, the dimension obtained by finding a linear projection of the neural population vector that distinguishes attack from sniffing periods (0.5% ± 0.1% of variance, N = 3 mice, p<0.001, Supplemental Figure S2B) Examining the activity of individual neurons that were weighted strongly in the integration dimension revealed that around 20% of neurons per animal showed ramping and persistent activity (Supplemental Figure S2E, F), with ramping neurons typically but not always preferentially activated by male intruders (Supplemental Figure S2D). Thus, the ramping dimension reflects a signal that is present at the level of at least some individual neurons, but is also an emergent property of the population (Ebitz and Hayden, 2021).

To understand what features (if any) the VMHvl integration dimension represented, we examined its correlation with the animals’ behaviors. We constructed a cumulative distribution function (cdf) of the level of activation along the integration dimension during three behaviors: sniffing, dominance mounting, and attack, in all imaged animals. We found that sniffing consistently occurs at low values of this dimension, dominance mount at intermediate values, and attack at high values (distribution mean, sniffing: 0.31, dominance mount: 0.67, attack: 0.81, N = 3 mice, Figure 2E). This suggests that during aggressive encounters, neural activity along this dimension grows as animals display escalating aggressive behaviors. Remarkably, periods of sniffing could be distinguished from attack or dominance mounting with high accuracy by simply thresholding this one-dimensional value (90.6%± 1.8%, N=3 mice, Figure 2D). The same method could also distinguish low- vs. high-intensity aggressive behavior (i.e., dominance mount vs. attack) at well above chance accuracy (67.3%± 2.9%, N=3 mice). Perhaps due to the slow dynamics of activity along this dimension, behaviors occurring close together in time, such as attack and sniff-attack (defined as periods of sniffing that occurred within one second prior to attack, as described by (Zhu et al., 2020)) could not be distinguished using this dimension. Remarkably, none of the other eight fit dimensions could be used to distinguish aggressive behaviors from sniffing with above chance accuracy (Figure 2F, G; Supplemental Figure S2C).

Because of its slow dynamics, activity in the integration dimension did not decay between individual bouts of behavior, and could not be used to predict the fine timing of actions. As a result, we could distinguish pauses (intervals of non-interaction) between sniffing bouts from pauses between attack bouts with high accuracy (83%± 2.1%, N=3 mice), whereas we could not distinguish sniffing bouts from the pauses between them (Figure 2D, Supplemental Figure S1A, right, Case 2). The level of activity along the integration dimension could not be predicted from the acceleration, facing angle, or velocity of the resident, or from the distance between the resident and intruder mouse, suggesting that the dimension did not simply encode pose-related features of the animal (mean: 0.35 ± 0.04 R^2^, n= 3 mice, Supplemental Figure S2A)

Thus, our unsupervised approach uncovered a one-dimensional signal in VMHvl^Esr1^ neural population activity that closely tracks and scales with an animal’s escalating level of aggressiveness and is reflected in the activity of approximately 20% of individual VMHvl^Esr1^ neurons. Different aggressive actions are associated with specific levels of activity along this dimension, suggesting an aggression-intensity code in VMHvl^Esr1^ activity. The fact that few VMHvl^Esr1^ neurons are tuned for specific aggressive actions suggests that this intensity code is translated into distinct actions by downstream circuits (Falkner et al., 2020). This correlation between VMHvl^Esr1^ activity and aggression gains additional causal support from the observation that increasing the intensity of optogenetic stimulation of VMHvl^Esr1^ neurons progressively evokes sniffing, dominance mounting and attack (Lee et al., 2014), actions that can be decoded from the integration dimension as its activity ramps up.

### VMHvl contains an approximate line attractor that integrates to create aggressive motivation

We next investigated how the fit linear dynamical systems contribute to the overall dynamics of population activity. For visualization purposes, we used principal component analysis (PCA) to find the first two principal components (PCs) of the dynamics of the fit rSLDS model (Figure 3A). In all imaged animals, PC1 showed slow ramping dynamics that escalated from low activity during the initial phase of social interaction when sniffing occurred, to higher activity as the animal displayed the first bouts of dominance mounting, to yet higher activity as the animal switched to attack behavior (Figure 3A, B, PC_1_ (behavior-triggered average, N = 3 mice)). Like the integration dimension in the full rSLDS space (Figure 2A), activity along PC1 remained persistently high throughout the attack period, suggesting that the integration dimension is a dominant component of activity in VMHvl. The component of neural activity along PC2 showed high activity when a new intruder was introduced but was otherwise low in all animals (Figure 3B, PC_2_ (behavior-triggered average, N = 3 mice)). Together these first two PCs accounted for 69.1%± 1.2% of the total variance in VMHvl activity across animals (N=3 mice).

**Figure 3:**
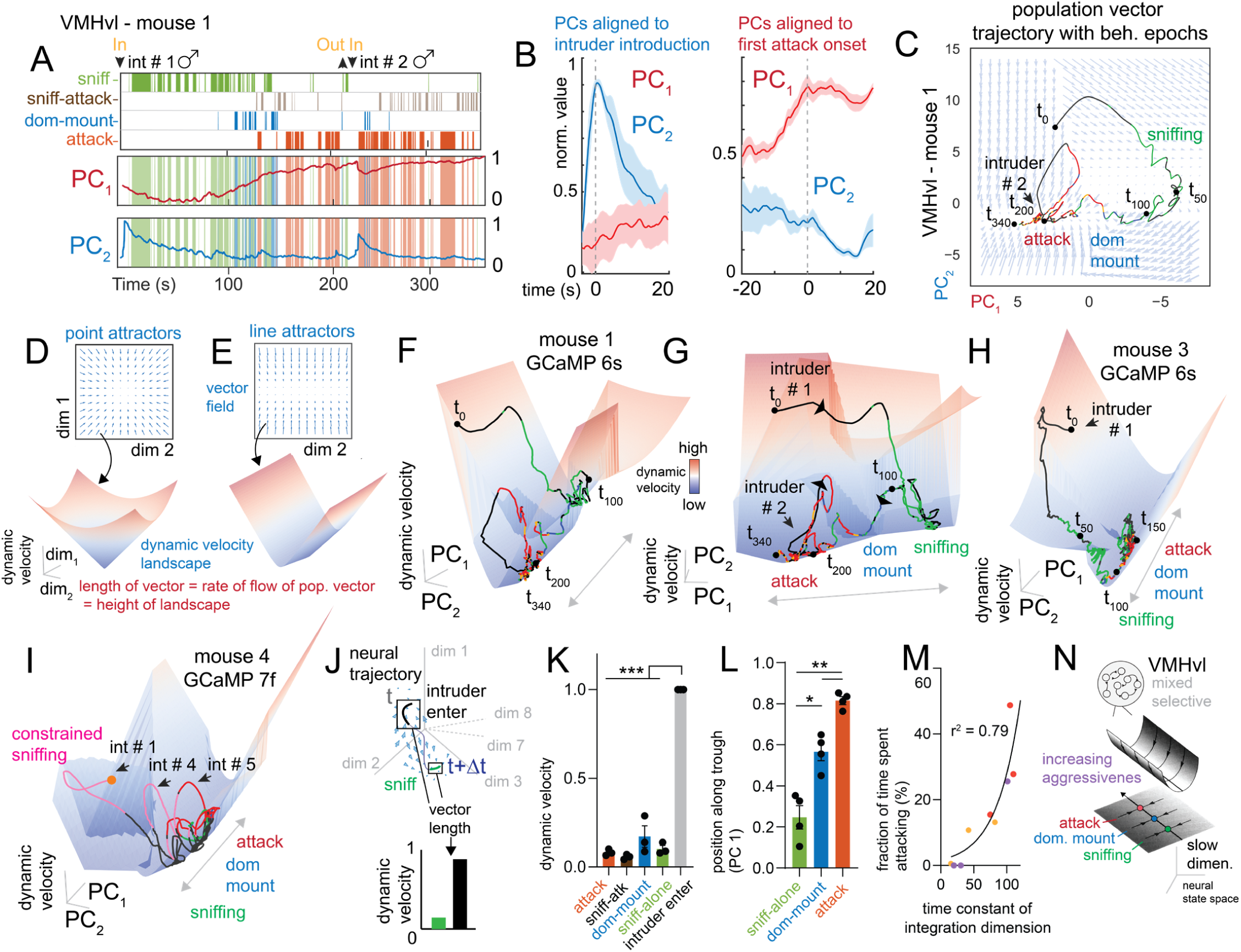
VMHvl contains an approximate line attractor that integrates to create aggressive motivation. A: state and behavior rasters shown with principal components of latent factors for example VMHvl mouse 1. B: behavior triggered average of top two principal components aligned to intruder introduction or first attack onset (n = 3 mice). C: neural state space with population trajectories for VMHvl mouse 1 colored by behaviors performed by the resident mouse. The inferred flow field of the model is shown behind trajectories. D,E: inferred dynamics shown as a dynamic velocity landscape, where the size (scalar value) of flow field vectors is represented as the height of the 3D landscape. F,G: inferred dynamic velocity landscape in VMHvl mouse 1. H: inferred dynamic velocity landscape for VMHvl mouse 3. I: inferred dynamic velocity landscape from a separate mouse where GCaMP 7f was used, showing a similar trough shaped landscape. J: calculation of dynamic velocity: the average vector length across dimensions for all time points of a given behavior. K: quantification of dynamic velocity (n = 3 mice, ***p<0.001). L: position of various behaviors along trough, i.e PC1 in neural state space (n = 4 mice, **p<0.005, *p<0.01). M: relationship between the time spent attacking and the time constant of the integration dimension (r^2^: 0.79, n = 9 animals; orange points are data from Karigo et al., 2021, purple points are data from Remedios et al., 2017, yellow points are data from Yang & Anderson, 2022). N: dynamical analysis finds an integration dimension that functions as an approximate line-attractor encoding an aggressive-intent drive in VMHvl.

By identifying the most-likely state of the fit rSLDS model at each point in neural activity space, we can create a flow field showing how we expect neural dynamics to evolve within the 2D space spanned by the first two PCs, in the absence of external input. Doing this revealed a region of low vector flow that forms an approximate line attractor, meaning that the state of the system tends to move towards points along the line in the absence of external input (Figure 3C, t_50_-t_340_). Line attractors are powerful coding mechanisms that allow neural populations to integrate inputs and robustly encode continuous variables such as head direction or reward value (Ganguli et al., 2008; Hulse and Jayaraman, 2020; Mante et al., 2013; Seung, 1996). The approximate line attractor we observed in VMHvl^Esr1^ activity is the result of the slow (integration) dimension of our dynamical system. Because all but one pattern of neural activity in the system have shorter time constants and decay away comparatively quickly, activity evoked by external sensory perturbations quickly settles into a one-dimensional, slowly decaying pattern. Continuous sensory or internal input during social interactions produce ongoing perturbation of the system state in neural activity space, causing it to gradually drift along the length of the attractor. As activity travels along the approximate line attractor, animals display a progression of increasingly aggressive behaviors, from sniffing to dominance mount to attack (Figure 3C, Supplementary Video 1).

To visualize better the dynamical landscape of the rSLDS model, we converted the 2D flow field into a 3D landscape, by converting the length of the vector at each position in neural state space (proportional to the rate of change of activity in the system at that point; which we term dynamic velocity; see *Methods*) into the height of the landscape (Figure 3D, E). In this representation, a line attractor appears as a trough or gully, whereas a point attractor would appear as a locus of stability at the base of a cone (Figure 3D, E, (Seung, 1996)). We observed a dynamic velocity landscape with a trough structure for every imaged animal (Figure 3F-I, Supplemental Figure S3): the short time constants of the non-integration dimensions produce the steep walls of the trough (corresponding to long flow-field arrows), whereas the long time constant of the integration dimension produces a long, flat gully (corresponding to short flow-field arrows), along which neural activity slowly evolves (Supplementary Video 2).

While the animals’ behavior was only strongly correlated with activation along the integration dimension, it is possible that the rate of change of dynamics in other dimensions might also show behavior-specific patterns. To test this possibility, we computed each behavior’s ‘dynamic velocity’: the average length of the vector field across all dimensions at all time points occupied by a behavior (Figure 3J). We found that time points associated with entry of the intruder mouse had the highest flows in all animals and were present on the walls of the trough, while all other behaviors resided within the base of the trough with little to no flow (Figure 3K). To determine where exactly behaviors resided along the axis of the trough, we quantified the average normalized value of PC1 for various behaviors across animals. This confirmed that evolution of the system along the trough corresponds with the animal displaying a progressive escalation of aggressive behaviors and was consistent with our earlier observation of an integration dimension in VMHvl dynamics (sniff-alone: 0.24± 0.05, dominance-mounting: 0.56 ± 0.04, attack: 0.81 ± 0.02, N =3 mice, Figure 3L).

### The time constant of the integration dimension in VMHvl predicts levels of aggressiveness across animals

While imaging from VMHvl in three different mice expressing GCaMP6s always revealed a single integration dimension with a long time constant, the magnitude of this time constant varied across animals. Strikingly, we observed a trend in which animals that displayed more aggressive behavior (calculated as the fraction of time spent attacking) also exhibited an integration dimension with a longer time constant (Figure 3M, red points). To validate our observation, we performed additional VMHvl^Esr1^ imaging in three more animals using the faster calcium indicator GCaMP7f; we again observed a single integration dimension in each of the three additional mice (Figure 3I, Figure 3M yellow points Supplemental Figure S3K-M). We also re-analyzed VMHvl^Esr1^ imaging from three additional animals from a previous dataset (Remedios et al., 2017) where animals displayed varying amounts of aggressiveness and where GCaMP6s was used as the calcium indicator (Figure 3M, purple points). Remarkably, all nine animals (6 recorded with GCaMP6s and 3 with GCaMP7f) exhibited a similar relationship between integration time constant and time spent attacking (Figure 3M, r ^2^ = 0.79, n = 9 animals). This striking correlation of integration time constant with time spent attacking suggests that aggressiveness may have a neural correlate in the intrinsic dynamics of VMHvl^Esr1^ neurons.

To summarize, our unsupervised dynamical systems analysis of Esr1^+^ neuronal population activity in VMHvl identified an approximate line attractor in neural state space, along which activity evolved with slow ramping dynamics and persistent activity (Figure 3N). Our analysis further suggests that this nucleus encodes more than just the sex of the intruder, perhaps reflecting an internal motive state of aggressiveness (Falkner et al., 2016; Falkner et al., 2020; Remedios et al., 2017).

### Mating behaviors are represented using rotational dynamics in the MPOA

Since rSLDS was able to discover evidence for integration in VMHvl, we next examined whether the same analysis would uncover population dynamics important for mating in MPOA, by fitting models to MPOA^Esr1^ neural data from interactions with female intruders (Karigo et al. 2021).

Fit models of MPOA required three rSLDS states in every animal, with USV+ mounting and intromission mostly occurring in single but different states (Supplementary Figure S4). Unlike in VMHvl, mating behaviors were closely aligned to the onset of individual neural states, and state bouts were similar to behavior bouts in their duration (Supplementary Figure S4 D, E). Upon examining the dynamics matrix associated with states where mating behaviors such as USV^+^ mounting (state 2) occurred in our fit models of MPOA^Esr1^ neurons, we did not find any evidence of dimensions with long time-constants (Figure 4A). Instead, the principal components of dynamics in the fit rSLDS model showed fast dynamics that were highly correlated with the occurrence of specific behaviors. Across all animals, PC1 of the fit dynamical system showed high activity during the introduction of a new female intruder, as well as at the onset of individual USV^+^ mounting bouts (Figure 4B, C, behavior triggered average, N = 3 mice), whereas PC2 showed modulation during intromission events (Figure 4C).

**Figure 4:**
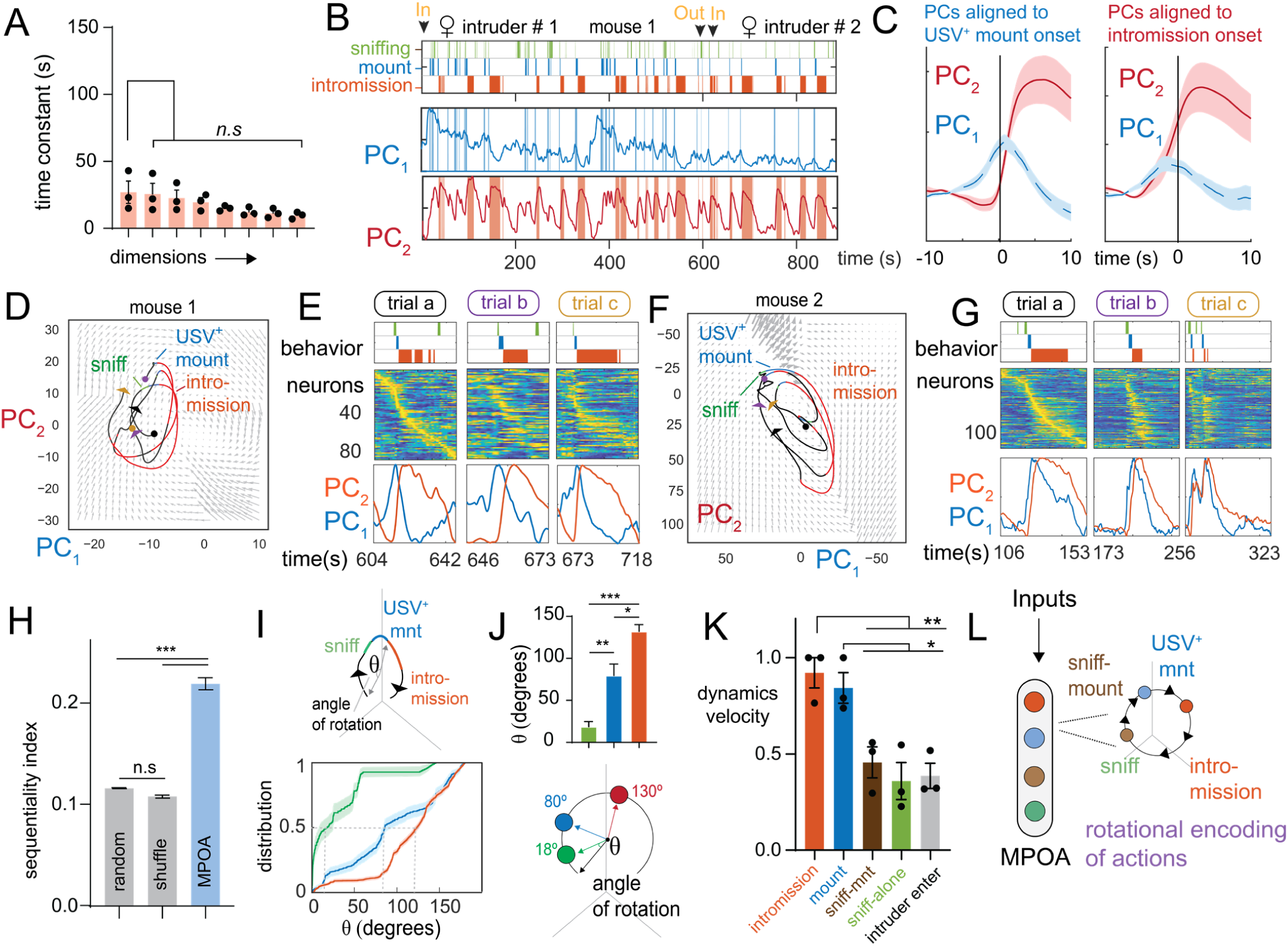
Mating behaviors are represented using rotational dynamics in the MPOA. A: time constants from dynamics matrix of mating behavior-enriched state in MPOA (n = 3 mice) B: state and behavior rasters shown with principal components of latent factors for example MPOA mouse 1. C: behavior triggered average of top two principal components aligned to USV+ mount onset and intromission onset (n = 3 mice). D: neural state space with rotational population trajectories from mating episodes shown in E of MPOA mouse 1, colored by behaviors performed by resident mouse. E: sequential activity of MPOA neurons during mating episodes whose rotational population trajectories are shown in D. F,G: same as D,E but for MPOA mouse 2. H: quantification of sequential index for MPOA (n = 3 mice, ***p<0.001). I: calculation of angle of rotation (θ) aligned to the start of sniffing during mating episodes (top). Empirical cumulative distribution of θ for various behaviors (n = mice, bottom). J: quantification of θ for various mating behaviors (n = 3 mice, ***p<0.001, **p<0.005, *p<0.01, top). Schematic depicting θ for mating behaviors (bottom). K: quantification of dynamic velocity for mating behavior in MPOA (n = 3 mice). L: dynamical analysis in MPOA find behaviorally tuned factors that displays a rotational encoding of actions.

The flow-field produced by the model visualized in 2D by PCA revealed that population activity vector dynamics was dominated by a rotational flow, with neural activity during a given trial undergoing a series of periodic orbits (Figure 4D, F). Moreover, the phase of flow rotation was tightly correlated with the animal’s progression through stages of reproductive behavior (sniffing, mounting, and intromission) (Figure 4I,J Supplemental Figure S5). Each rotational trajectory corresponded to the sequential activation of different neurons during different stages of mating (Figure 4E,G, Supplemental Figure S5). This sequential activity was not an artifact of sorting neural activity, as the sequentiality of the data (seq. index = 0.22 ± 0.01, N = 3 mice, as defined in Zhou et al., 2020) was significantly greater than shuffled data or random matrices of similar sizes (shuffle seq. index = 0.10± 0.002, N = 3 mice Figure 4H)

We assessed the relationship between the phase of rotational trajectories and behavior by calculating the angle of rotation aligned to the start of sniffing during mating episodes (Figure 4I). This revealed that the phase of rotation tracked the sequential events during a mating cycle, with sniffing, mounting and intromission defined by characteristic angles of rotation (sniffing: 18.6° ± 6.2 °, mounting: 79.61°±13.6 °, intromission: 132.2°± 8.1 °, N = 3 mice Figure 4J). Importantly, rotational dynamics during mating events were seen in each of the three animals for which we fitted rSLDS models (Supplemental Figure S5). The dynamic velocity associated with behaviors in MPOA revealed high flows associated with consummatory behaviors, which was markedly different from the slow dynamics during aggressive behavior in VMHvl (Figure 3L, 4K).

Thus, unlike the slow ramping dynamics identified in VMHvl, rSLDS discovered faster, behaviorally time-locked rotational dynamics in MPOA. Furthermore, we found that during mating bouts, MPOA exhibits sequential activation of a series of neurons, whose firing is aligned with distinct stages of the behavior (Figure 4L).

### Distinct neural coding schemes for similar behavior in VMHvl vs MPOA

Our dynamical systems analysis of social behavior in VMHvl and MPOA identified two distinct neural coding schemes for aggression vs mating behavior in the two nuclei. The integration dimension found in VMHvl is strikingly absent in MPOA, emphasizing the slow ramping nature of dynamics in VMHvl compared to the fast behaviorally aligned dynamics in MPOA (Figure 5A, B). Behaviors such as attack occur during periods with relatively small rates of change in VMHvl population dynamics, and the system resides within the trough of a dynamic velocity landscape. In contrast, consummatory behaviors in MPO have high rates of change in population dynamics, due to the rotational nature of the system (Figure 5C, D). Yet both regions possess a signal that tracks aggressive and mating behavior: the value of the integration dimension in VMHvl is correlated with escalating aggressive behavior, while the angle of rotational dynamics in MPOA is correlated with progression of mating episodes (Figure 5E, F). The rotational dynamics in MPOA are accompanied by sequential activation of individual neurons, while sequential activity is absent during aggression episodes in VMHvl (Figure 5G, H). Thus the same method, rSLDS, uncovers two distinct forms of dynamics for encoding similar social behavior in VMHvl and MPOA (Figure 5I, L).

**Figure 5:**
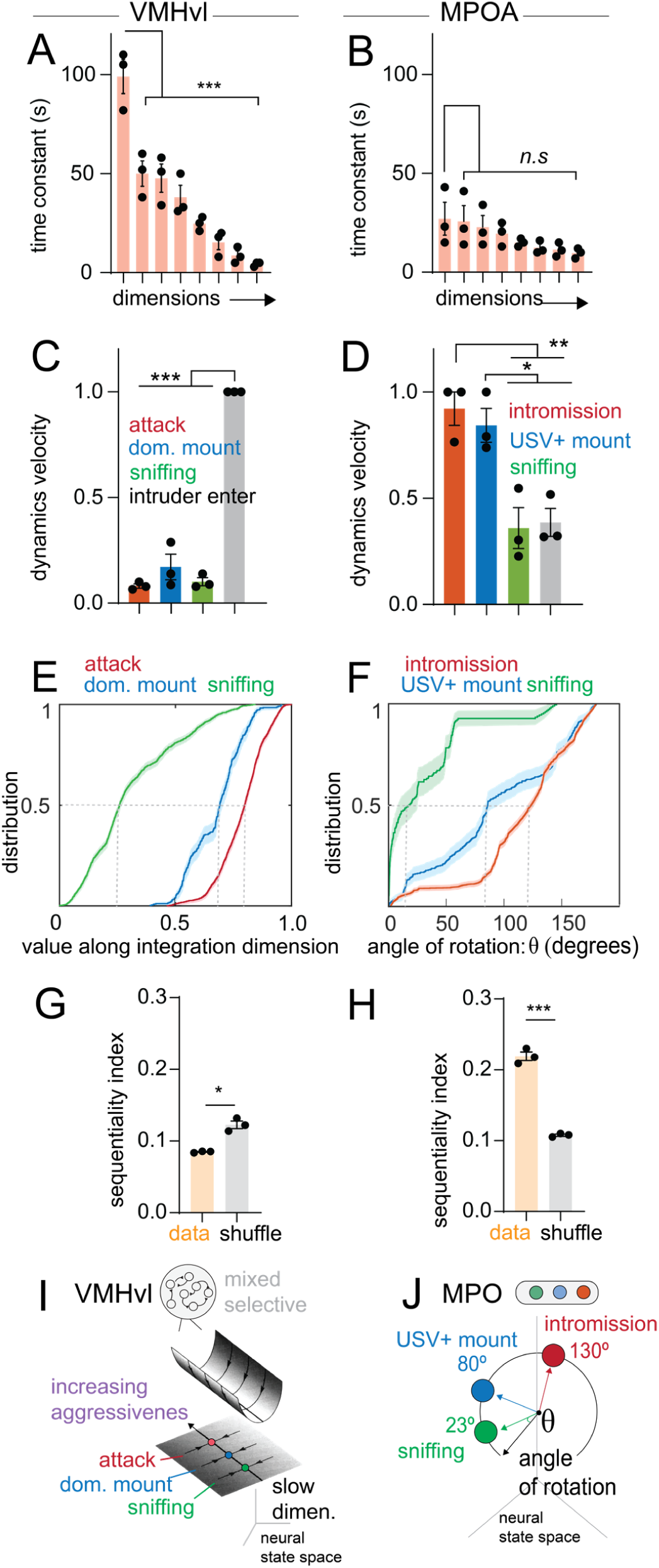
Distinct neural coding schemes for similar behavior in VMHvl vs MPOA. A: time constants from dynamics matrix of attack behavior-enriched state in VMHvl (n = 3 mice), reproduced from Figure 2C. B: time constants from dynamics matrix of mating behavior-enriched state in MPOA (n = 3 mice), reproduced from Figure 4A. C,D: quantification of dynamic velocity in VMHvl (C) and MPOA (D). Reproduced from Figure 3K and Figure 4K resp. E: empirical cumulative distribution of value of integration dimension (normalized) in VMHvl for various behaviors, reproduced from Figure 2E. F: empirical cumulative distribution of angle of rotation (normalized) in MPOA for various behaviors, reproduced from Figure 4I. G: Sequentiality index in MPOA (n = 3 mice), reproduced from Figure 4E. H: Sequentiality index in VMHvl in aggression (n = 3 mice). I: dynamical analysis finds an integration dimension that functions as an approximate line-attractor encoding an aggressive-intent drive in VMHvl. J: dynamical analysis in MPOA find behaviorally tuned factors that displays a rotational encoding of actions.

### VMHvl also contains an integrator dimension encoding reproductive state

Our discovery of line attractor dynamics for aggression in VMHvl and rotational dynamics for mating in MPOA raised an important question: do these contrasting dynamics reflect fundamental properties of aggressive vs. mating behavior, or do they reflect differences in the functional roles of the two nuclei with respect to social behaviors? To address this, we fit models to neural activity from VMHvl^Esr1^ neurons during social interactions with females.

Fit models required three rSLDS states in all animals, with mating behaviors (USV+ mounting and intromission) mostly present in one state (state 3, Supplementary Figure S6A-H). As in the case of aggression, bouts of the mating-dominant state in VMHvl lasted much longer than individual bouts of mating behavior (Supplementary Figure S6D, H). Given the persistence of the mating states discovered by rSLDS in VMHvl, we examined the time constants associated with each dimension of the mating-dominant state in VMHvl (state 3, Supplementary Figure S6). As for aggression, we found a single integration dimension with ramping and persistent activity, with a large time constant in every animal (Figure 6A red dot, 6B). We next asked if we could distinguish mating vs non-mating behavior from the value of this single dimension. Surprisingly, we could distinguish pairs of behaviors with high accuracy (intromission vs sniffing: 0.94± 0.02 N = 2 animals (third animal did not display intromission), USV^+^ mounting vs sniffing: 0.78± 0.04 N = 3 animals Supplementary Figure S6I). As for aggression, we could distinguish periods of non-interaction between mounting bouts from those between sniffing bouts (Supplementary Figure S6I), but could not distinguish sniff-mount (sniffing occurring within one second of mounting) from mounting. Finally, the cumulative distribution of the value of the integration dimension during sniffing, mounting, or intromission revealed that sniffing occurs at the lowest values of this dimension, USV^+^ mounting occurs at intermediate values, while intromission occurs at the highest values of this dimension (Figure 6C). All of these analyses indicate that the dynamics of VMHvl during reproductive behavior closely resemble those observed in VMHvl during aggression. To continue our comparison, we projected our high dimensional model into a two-dimensional space using PCA (Figure 6D). The resulting projections appeared similar to those seen in VMHvl during aggressive encounters with males. PC1 had low activity during sniffing bouts, with activity ramping up as early USV^+^ mounting (typically not proceeding to intromission) was displayed, and remaining persistently high for hundreds of seconds as the animal showed several cycles that progressed from sniffing to mounting to intromission (Figure 6D). Conversely, PC2 showed high activity during the introduction of a new intruder but was otherwise low. Examining the underlying vector field revealed regions of stability in mating-related states that resembled an approximate line attractor and a trough shaped dynamic velocity landscape, with neural activity moving along the line attractor as the animal progressed from the appetitive to consummatory phases of mating (Figure 6E, F); apparent rotations in Figure 6E occurred only when introduction of a new intruder transiently increased activity along PC2. Across animals, dynamic velocity analysis revealed that only times of intruder introduction had high flows, while all other behaviors had little flow and were present in the trough of the dynamic velocity landscape (Figure 6G).

**Figure 6:**
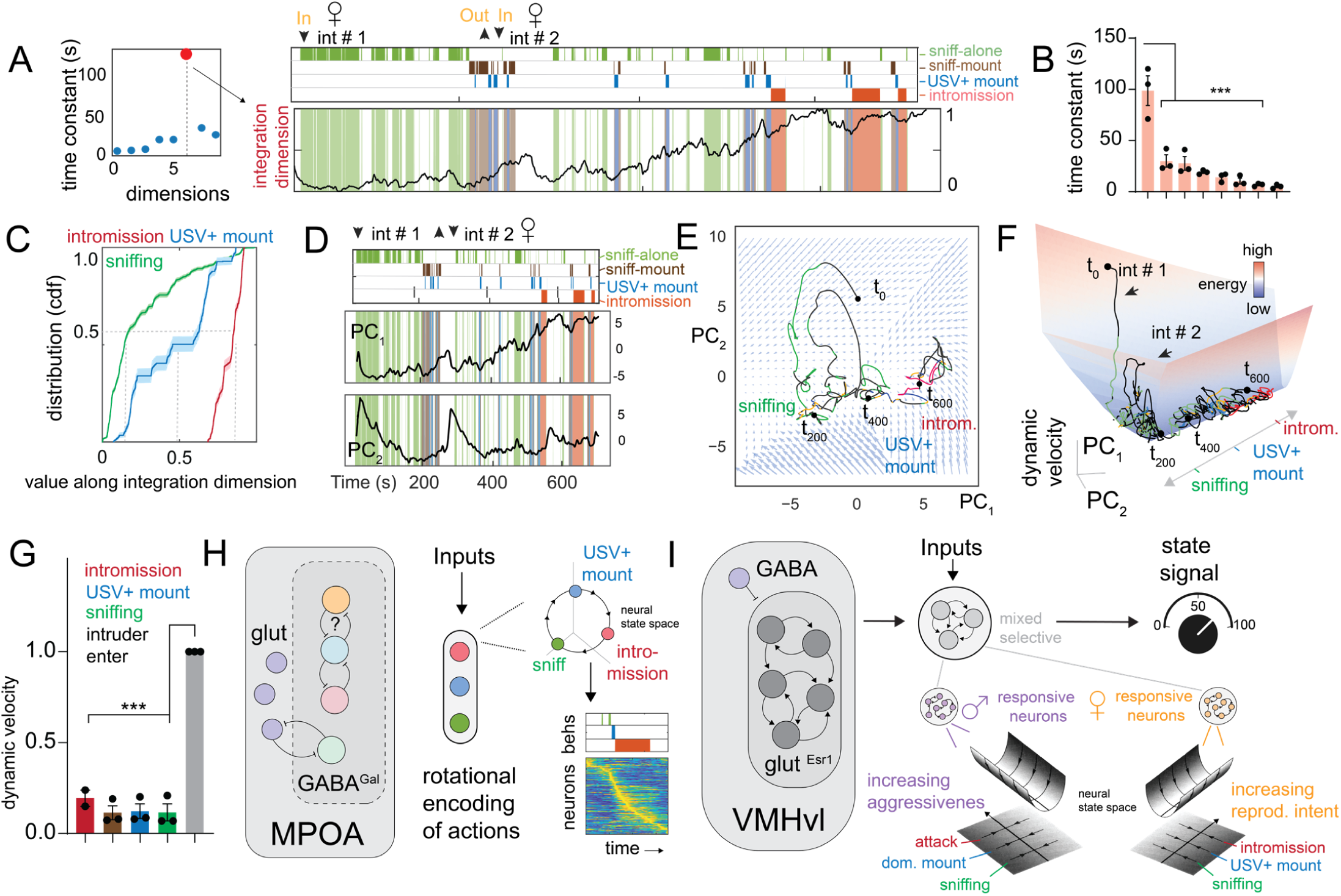
VMHvl contains a separate approximate line attractor encoding reproductive intent. A: time constants of dynamics matrix of mating enriched state from example VMHvl mouse 1. The red dot highlights the largest time constant or integration dimension. B: Time constant from all animals arranged in decreasing order. The integration dimension (dim 1) is significantly larger than all other dimension (p < 0.001, n = 3 mice). C: empirical cumulative distribution of value of integration dimension (normalized) for various behaviors. D: state and behavior rasters shown with principal components of latent factors for example VMHvl mouse 1. E: neural state space with population trajectories for VMHvl mouse 1 colored by behaviors performed by the resident mouse. The inferred flow field of the model is shown behind trajectories. F: dynamic velocity landscape with population trajectory colors by behavior. G: quantification of dynamic velocity during mating behavior in VMHvl across animals (p<0.001, n = 3 mice). H: MPOA contains distinct cell populations that are organized by rotational dynamics and encodes actions. I: VMHvl contains mixed selective neurons that are organized by line attractor dynamics that encode a representation of escalating motivational states.

Lastly, examining the weighted activity of neurons that contribute to the integration dimension revealed several cells with ramping and persistent activity during male-female encounters (Supplementary Figure S6J). The integration dimension seen during mating behavior was biased towards neurons tuned to female intruders, and was largely non-overlapping with neurons that created the aggressiveness integration dimension seen in interactions with males (Supplementary Figure S6F).

Thus, the application of rSLDS to data recorded during male-female social interactions uncovered a representation of mating in VMHvl in which activity evolves along a line attractor, possibly reflecting the intensity of reproductive drive. This attractor is similar to that found in VMHvl during aggression, but incorporates primarily female-selective rather than male-selective neurons (Figure 6H, I). In contrast, behavior-locked rotational dynamics like those found in MPOA during mating behavior were not observed in VMHvl. We therefore conclude that line-attractor dynamics are a general feature of VMHvl, rather than a signature of a particular type of social behavior.

## Discussion

The dramatic and specific behavioral phenotypes obtained by optogenetically activating genetically distinct hypothalamic neuronal cell classes has supported the idea that the hypothalamus controls innate behaviors via dedicated, behavior-specific neural subsets. This conclusion has also been drawn from c-fos labeling experiments performed in combination with transcriptomic identification of hypothalamic cell types in the pre-optic area (POA), which has revealed neuronal subtypes selectively activated during parental, reproductive or aggressive behaviors (Moffitt et al., 2018). In MPOA, moreover, different subsets of neurons have been shown to project to distinct downstream targets, with minimal collateralization (Kohl et al., 2018). Together, these data have reinforced the prevailing view that hypothalamic nuclei are comprised of genetically specified, behavior-specific neuronal subpopulations connected in developmentally determined pathways or “labeled lines” (Ishii et al., 2017; Kohl et al., 2018).

It is tempting to generalize these findings to other hypothalamic nuclei that control different survival behaviors. Indeed, studies of the arcuate nucleus (ARC) have revealed genetically distinct subpopulations that control feeding behavior via non-collateralizing projections to different targets (Sternson, 2013). Like MPOA, moreover, ARC is predominantly GABAergic. However, other behaviorally relevant hypothalamic nuclei exhibit very different neurochemical, cytoarchitectonic and connectional features. For example, VMHvl and PMv, which play a dominant role in the control of aggression (Lee et al., 2014; Lin et al., 2011; Stagkourakis et al., 2018; Yang et al., 2013), are predominantly glutamatergic, with a surrounding GABAergic “shell.” Furthermore, VMHvl^Esr1^ neurons collateralize extensively to their downstream targets (Lo et al., 2019), unlike MPOA and ARC (Figure 1B). Finally, unlike the case in the POA (Moffitt et al., 2018), immediate early gene expression and scRNAseq analysis in VMHvl has failed to reveal a robust relationship between transcriptomic cell type identify and behavior-specific activation (Kim et al., 2019). The functional significance of these differences between MPOA and VMHvl has until now remained unclear.

### MPOA and VMHvl encode social behaviors in a digital vs. analog manner, respectively

Here we report that MPOA^Esr1^ and VMHvl^Esr1^ neurons utilize very different schemes for the neural coding of behavior, despite the fact that optogenetic stimulation of these neurons evokes mating and aggression, respectively, two highly related social behaviors. GCaMP imaging of VMHvl^Esr1^ neurons during social interactions has revealed very few cells tuned to specific behavioral actions, unlike the case in MPOA in which many neurons are only active during a specific preferred behavior (Karigo et al., 2021; Remedios et al., 2017). In contrast, we show that specific aggressive behaviors can be decoded from features of population neural activity in VMHvl. Thus, MPOA represents behavior via a cell identity code, while VMHvl does so via a population code.

Our studies here reveal the nature of this population code. rSLDS analysis of VMHvl neural activity during male-male social interactions, projected by rSLDS into an 8-dimensional latent factor space, revealed one dimension with a long time-constant that exhibits progressively increasing activity during escalating aggressive encounters. Different aggressive actions, such as dominance mounting or attack, can be distinguished with high accuracy by the neural activity along this single dimension. In a topological representation, these dynamics can be visualized as a progression along a stable “trough” or gully, which has the characteristics of a line attractor. In contrast, a similar analysis of MPOA revealed rotational dynamics, in which the population cycles through distinct states generated by the activity of behavior-specific cell types during each bout of mating. Put simply, VMHvl coding of behavior appears to be analog, while MPOA coding of behavior appears more digital.

The different neural codes for behavior we have uncovered in VMHvl and MPOA fit well with their distinct neurochemical and cytoarchitectonic features. Glutamatergic neurons in VMHdm exhibit local connectivity (Kennedy et al., 2020) which, if true for VMHvl as well, can in theory support the persistent activity necessary for the ramping activity and attractor dynamics we observe. Conversely, the fact that the majority (85%) of MPOA neurons (including those that control mating) are GABAergic could provide a substrate for reciprocal inhibitory connections between action-specific subpopulations. Such connectivity could produce winner-take-all dynamics or feed-forward dis-inhibitory circuits that control transitions between sequential action phases of mating (e.g., from mounting to intromission), giving rise to the rotational dynamics observed in neural data. The existence of such circuits in MPOA can be investigated using slice physiology or in vivo imaging experiments once appropriate cell type-specific markers become available.

Why should MPOA and VMHvl utilize such different strategies for the coding of closely related social behaviors? It is tempting to attribute this difference in population dynamics to distinct features of reproductive vs aggressive behavior. For example, aggressive encounters can dynamically escalate or de-escalate, in order to avoid serious injury or death to the combatants, whereas male mating typically proceeds to completion (ejaculation) once initiated. These differences are well-suited to control by ramping and cycling neuronal dynamics, respectively. In this view, the different properties and coding strategies of VMHvl and MPOA may have evolved to be optimally adaptive for fighting and mating, respectively.

However, our analysis also revealed line attractor dynamics in the subset of VMHvl^Esr1^ neurons that is female-tuned and active during mating (Remedios et al., 2017). This suggests that line attractor dynamics are a general property of behavioral coding by VMHvl, not an aggression-specific feature. By the same token, MPOA contains specific neurons highly tuned to attack, in contrast to VMHvl (although it is not yet clear whether these neurons play an aggression-promoting or –inhibiting role). Thus, our analysis suggests that MPOA and VMHvl more likely encode different aspects of mating behavior, such as action selection vs. drive state intensity, respectively. A generalization of this view predicts that the hypothalamus should contain at least one additional nucleus with MPOA-like behavioral coding, which controls aggression. Indeed, the anterior hypothalamic nucleus (AHN), which has a similar neurochemical and cytoarchitectonic structure as MPOA, can promote defensive attack (Nelson and Trainor, 2007; Xie et al., 2022); it will be interesting to see whether rotational dynamics are observed in this structure. Conversely, this view predicts that PMv, which promotes aggression and is primarily glutamatergic like VMHvl (Stagkourakis et al., 2018), may utilize population coding of behavior and exhibit line attractor dynamics.

### Potential functions of the VMHvl line attractor

Line attractors have been identified in cortical and hippocampal regions involved in cognitive functions, such as decision-making, spatial mapping and sensory discrimination. It is unexpected to find such neural dynamics in the hypothalamus, which is widely viewed as controlling innate behaviors via action-specific cell types similar to those observed in MPOA. What function(s) might such attractor dynamics serve, in the context of innate behaviors? Two explanations are possible, which are not mutually exclusive.

First, progression along the line attractor may encode the intensity of an internal motive state of aggressiveness. This is supported by our finding that the integration dimension that contributes to this attractor can distinguish periods of non-social interaction during high-vs. low-intensity phases of aggressive escalation (Figure 2D). In this view, the line attractor functions to maintain the system in a stable motive state, during variable behavior. Note that this line attractor traverses the boundary between two of the linear dynamic states identified by rSLDS (Figure 3C). Therefore, the brain state of aggressiveness is a property of the overall dynamics of VMHvl population activity, not just of one rSLDS “state”. The idea that an internal emotion or motive state may be encoded by a continuous line attractor provides a way to stably maintain such a state, whether or not its associated behaviors are overtly expressed on a moment-to-moment basis. The neural mechanisms that underly this state could include recurrent excitatory networks, neuromodulatory systems or a combination of both, as observed in VMHdm neurons controlling a fear-like state (Kennedy et al., 2020).

An alternative explanation is that the line attractor may serve as an integrator that accumulates “evidence” used to make behavioral decisions, such as the decision to switch from sniff to dominance mount, or from dominance mount to attack. Such a function would require that different behaviors be triggered at different threshold values of the integrator. This type of ramp-to-threshold mechanism has been suggested to control sequential actions during male courtship behavior in *Drosophila* (McKellar et al., 2019) and predator escape in mice (Evans et al., 2018). These two functions are not incompatible: the attractor could encode both the intensity of an internal state, and (indirectly) the selection of actions at different state intensities.

Finally, line attractor dynamics could serve useful functions in the context of behavioral plasticity and individual variation. For example, VMHvl^Esr1^ neurons show increased selective tuning for male vs. female intruders as a function of social experience (Remedios et al., 2017), and exhibit a form of long-term potentiation that underlies the increase in aggressiveness that occurs when mice win a series of fights (Stagkourakis et al., 2020). It will be interesting to determine whether changes in flow field dynamics or attractor properties are associated with these forms of experience-dependent plasticity. Finally, we note that differences in line attractor properties were observed among mice which exhibited different and characteristic levels of aggressiveness (Figure 3M). It is possible that individual differences in aggressiveness may reflect, or be caused by, individual constraints on population dynamics in VMHvl.

### Computational vs. behavioral measurements of internal brain states

Traditional approaches to identifying internal drive or motivational states have relied on learned behavioral tests such as operant conditioning. Neural activity that correlates with the execution and intensity of the conditioned instrumental behavior (e.g., bar-pressing), as determined using Progressive Ratio experimental designs, is interpreted as encoding the motivational state and its relative strength (Atasoy et al., 2012). While this approach has worked well for homeostatic motivational states such as hunger and thirst, there are several problems with generalizing it to innate social behaviors. First, it cannot be applied while the animal is directly engaged in a behavior such as aggression, because it requires the animal to cease directing its behavior towards the conspecific and to re-direct it towards the conditioning apparatus (nose-poke or bar-press). Therefore, it can be used to measure an animal’s motivational drive to seek out the opportunity to engage in a social behavior (Covington et al., 2019; Falkner et al., 2016; Golden et al., 2019), but not its state during engagement in the social behavior *per se*. Second, it substitutes the performance of a learned, artificial (instrumental) behavior for that of an innate behavior. Finally, it restricts the analysis of neural correlates of the motivational state to recordings performed occurs during the performance of the instrumental behavior.

Here, we have identified neural correlates of aggressive and mating internal drive states by examining the dynamic landscape of population neuronal activity in VMHvl. This approach does not require any instrumental operant conditioning, but can be performed on data recorded during the naturalistic behavior itself. Moreover, the approach identifies the states purely from analysis of neural population activity, without any reference to behavioral annotation. Importantly, unlike neural correlates of operant conditioning which are observed during performance of the instrumental behavior (Falkner et al., 2016), the signal discovered by rSLDS analysis can be observed both when the animal is behaving, and when it is not behaving (i.e., during inter-bout pauses). Indeed, it is the persistence of this signal throughout both active attack behavior and intermittent periods of inactivity between attack bouts, that suggests it encodes an internal state rather than behavior itself (Figure S1A). That this persistence is not due to slow GCaMP dynamics is suggested by the fact that 1) it is not observed in MPOA; and 2) it is also observed using a faster version of GCaMP (7f).

Our conclusion that VMHvl^Esr1^ neurons in males encode an internal motive or drive state underlying aggression, rather than simply the sex identity of a conspecific male, is consistent with recent studies of this neuronal population in female mice. Unlike male mice, which only attack other males, lactating female mice attack intruders of both sexes. A subset of VMHvl^Esr1^ neurons in females that express the GPCR gene *Npy2r*, called ß cells, are both necessary for maternal aggression and sufficient to promote attack in non-aggressive virgin females (Liu et al., 2022). Bulk calcium measurements in behaving animals show that ß cells are equally active during maternal aggression towards both male and female intruders. However, these cells display lower activity in individual mice that are non-aggressive (Liu et al., 2022). These data suggest that VMHvl^Esr1^ ß cells in females encode aggressiveness rather than intruder sex; the same may be true for males.

### Testable predictions of the line-attractor model

Our rSLDS model of VMHvl dynamics makes several testable predictions and raises several interesting questions for future investigation. First, it predicts that once in the attractor, the system will return quickly to it following perturbations that move it out of this stable trough. This behavior is suggested by the brief excursion out of the attractor that occurs when a first intruder is removed and replaced by a second one. However, it would be ideal to demonstrate this directly by transiently activating neurons that contribute to the attractor, and determining whether the system rapidly returns to it following stimulus offset, as has been demonstrated for point attractors underlying working memory in ALM (Inagaki et al., 2019). Another prediction is that selectively inactivating the VMHvl^Esr1^ neurons that exhibit slow dynamics should eliminate the line attractor. Such experiments will require combined optogenetic perturbations and calcium imaging in this deep subcortical structure.

Single-cell RNAseq experiments have shown that the Esr1^+^ population in VMHvl contains 6-7 distinct transcriptomic subtypes (Kim et al., 2019). An interesting question is whether all, or just a subset, of these cell types contribute to attractor dynamics. This question can be addressed once genetic drivers that are specific for these subtypes are generated. An additional question is whether the slow dynamics observed for some VMHvl^Esr1^ neurons reflects recurrent connectivity between them, as has been demonstrated for fear-encoding neurons in VMHdm (Kennedy et al., 2020), or the release of slow-acting neuromodulators such as neuropeptides. Recurrent connectivity in VMHvl can be investigated by slice electrophysiology and ultimately by EM connectomics. VMHvl^Esr1^ neurons are known to express multiple neuropeptides, as well as receptors for neuropeptides and other neuromodulators. New sensors for detecting neuromodulator release (Sabatini and Tian, 2020; Sun et al., 2018), as well as methods for selectively perturbing neuromodulator function while simultaneously imaging population neural activity *in vivo*, should help to address these questions in the future.

### Evolutionary implications

The observation that neural activity in MPOA during social behavior reflects distinct subpopulations, whose activity is time-locked to specific behavioral actions, is consonant with the prevailing view that the hypothalamus encodes innate behaviors via developmentally determined cell types with hard-wired connectivity (Ishii et al., 2017; Kohl et al., 2018; Moffitt et al., 2018). In contrast, the mixed selectivity exhibited by VMHvl neurons is more often observed in the cortex (Fusi et al., 2016) where attractor dynamics have also been identified as a component of population coding. As in the cortex, coding in VMHvl is mediated predominantly by glutamatergic neurons, with GABAergic neurons in the shell apparently playing a more peripheral role, whereas in the MPOA behavioral control is carried out predominantly by GABAergic projection neurons (Chen et al., 2021; Karigo et al., 2021), similar to the basal ganglia (Graybiel, 2000; Klaus et al., 2019; Park et al., 2020). As the hypothalamus emerged in vertebrate evolution long before the cortex, it is tempting to speculate that VMHvl and related glutamatergic nuclei might have functioned like a primitive cortex, while MPOA and related GABAergic nuclei might have functioned analogously to the basal ganglia. Comparative studies of hypothalamic neural population dynamics in non-traditional model organisms may help to shed light on this hypothesis.

## Acknowledgments

We thank S. Ganguli and M. Schnitzer for discussions on conceptualizing this project at its inception. We thank H. Inagaki and L.F. Abbott for critical feedback on this manuscript, C. Chiu for laboratory management and G. Mancuso for administrative assistance, and members of the Anderson and Kennedy labs for helpful comments on this project. A. N. is supported by a National Science Scholarship from the Agency of Science, Technology and Research, Singapore. D.J.A. is an Investigator of the Howard Hughes Medical Institute. A.K. is supported by NIH R00MH117264. The content is solely the responsibility of the authors and does not necessarily represent the official views of the National Institutes of Health.

## Author contributions

D.J.A, A.K and A.N conceived of the project and wrote the manuscript, with input from T.K, B.Y and S.L. A.K performed the clustering analysis of single neurons and A.N performed all dynamical system modelling.

## Methods

### Data

#### Neural imaging data (Karigo et al., 2021, Remedios et al., 2017, Yang and Anderson, 2022)

We analyzed data from three sets of previous experiments (Karigo et al., 2021; Remedios et al., 2017; Yang and Anderson, 2022) All experiments were approved by the Institute Animal Care and Use Committee (IACUC) and the Institute Biosafety Committee (IBC) at the California Institute of Technology (Caltech). Briefly, for data obtained from Karigo et al., 2021, *Esr1^cre^* knock-in mice, in which GCaMP6s was expressed selectively in *Esr1* neurons in either the medial preoptic area (MPOA) or the ventrolateral subdivision of the ventromedial hypothalamus (VMHvl), were allowed to interact with BALB/c male and female intruders in a standard resident intruder assay (cite). Male or female intruders were introduced into the home case in a random order, with a 5-10 min interval between intruder session. Each session typically lasted 10-20 minutes. Behavior videos of interacting animals were annotated using a custom MATLAB-based interface. A total of 7 behaviors including sniffing, dominance-mount, attack, mount, intromission, interact (periods where animals were close to each other but other behaviors were absent) were annotated with male and female intruders. A head-mounted micro-endoscope was used to acquire Ca^2+^ imaging data at 15Hz from either MPOA^Esr1^ neurons (total of 583 neurons from 3 mice) or VMHvl^Esr1^ neurons (total of 421 neurons from 3 mice) for neural data analysis described in sections below.

For data obtained from Yang and Anderson, 2022, briefly, *Esr1^cre^* knock-in mice, in which GCaMP7f was expressed selectively in *Esr1* neurons in VMHvl, were allowed to interact with BALB/c male intruders in a standard resident intruder assay. In addition to the behaviors annotated for above, male intruders were also “dangled”, where the ano-genital region of the dangled intruder is held next to the resident mouse. A head-mounting micro-endoscope was used to acquire Ca^2+^ imaging data at 30Hz from VMHvl^Esr1^ neurons (386 neurons from 3 mice) for neural data analysis described in sections below.

For data obtained from Remedios et al, 2017, briefly, *Esr1^cre^* knock-in mice, in which GCaMP6s was expressed selectively in *Esr1* neurons in VMHvl, were allowed to interact with BALB/c male intruders in a standard resident intruder assay. A head-mounting micro-endoscope was used to acquire Ca^2+^ imaging data at 30Hz from VMHvl^Esr1^ neurons (358 neurons from 3 mice) for neural data analysis described in sections below.

### Neural data analysis

#### Tuning rasters for single neurons

We examined the tuning properties of single neurons in VMHvl^Esr1^ or MPOA^Esr1^ by creating behavior tuning rasters (Figure 1 C, D). We first computed the mean activity of each neuron for each the 14 manually annotated behavioral actions. To group neurons, we created a set of 40 regressors representing combinations of behavioral actions, and grouped neurons by which single regressor captured the most variance in each cell’s activity. In addition to regressors for individual behaviors, example regressors include signals such as all male-directed actions, all female-directed actions, all male-directed/female-directed/sex-invariant investigative behaviors, and all male-directed/female-directed/sex-invariant consummatory behaviors. Neurons for which no single regressor captured at least 50% of variance in behavior-averaged activity were omitted from the visualization (approximately 5% of cells.)

#### Computation of pose features for input to dynamical model

As external input to the dynamical model (see next section), we selected two features of animal pose estimates produced by the Mouse Action Recognition System (MARS, Segalin et al., 2020) The first of these is the distance between animals, computed as the distance between centroids of ellipses fit to the poses of the two mice. The second is the facing angle of the resident towards intruder mouse, defined as the angle between a vector connecting the centroids of the two mice and a vector from the centroid to the nose of the resident mouse.

#### Dynamical system models of neural data

We model neural activity using a recurrent switching linear dynamical systems (rSLDS) according to previous methods (Linderman et al, 2019, Linderman et al., 2017). Briefly, rSLDS is a generative model that breaks down non-linear time series data into sequences of linear dynamical modes. The model relates three sets of variables: a set of discrete states (z), a set of continuous latent factors (x) that captures the low-dimensional nature of neural activity, and the activity of recorded neurons (y). The model also allows for external inputs (u), for which we used the two pose features described in the previous section. The between these various levels is shown graphically in Supplementary Figure 13 and is described below. The model also allows for external inputs (u) which consists of extracted pose features including the distance between animals and the facing angle between the resident and intruder mouse.

The model is formulated as follows: At each time t = 1,2,…T_n_, there is a discrete state *z_t_* ∈ {1,2, …, *K* }. In a standard SLDS, these states follow Markovian dynamics, however rSLDS allows for the transitions between states to depend recurrently on the continuous latent factors (x) and external inputs (u) as follows:

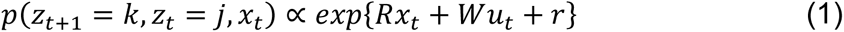

where *R, W* and *r* parameterizes a map from the previous discrete state, continuous state and external inputs using a softmax link function to a distribution over the next discrete states.

The discrete state *z_t_* determines the linear dynamical system used to generate the continuous latent factors at any time t:

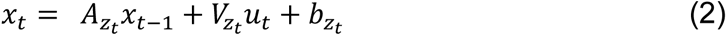

where *A_k_* ∈ ℝ*^d×d^* is a dynamics matrix and *b_k_* ∈ ℝ*^d^* is a bias vector, where *d* is the dimensionality of the latent space. Thus, the discrete state specifies a set of linear dynamical system parameters and specify which dynamics to use when updating the continuous latent factors.

Lastly, we can recover the activity of recorded neurons by modelling activity as a linear noisy Gaussian observation *y_t_* ∈ ℝ*^N^* where *N* is the number of recorded neurons:

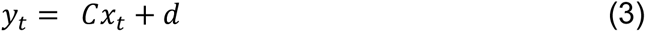

For *C* ∈ ℝ*^N×D^* and *d ∼ N*(0, *S*), a gaussian random variable. Overall, the system parameters that rSLDS needs to learn consists of the state transition dynamics, library of linear dynamical system matrices and neuron-specific emission parameters, which we write as:

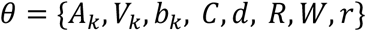

These parameters are estimated using maximum likelihood using approximate variational inference methods as described in detail in Linderman et al., 2019.

Model performance is reported as the *evidence lower bound (ELBO)* which is equivalent to the Kullback-Leibler divergence between the approximate and true posterior, *KL*(*q(x,z;φ*) || (*p(x,z|y;θ)*) using 5-fold cross validation. Code used to fit rSLDS on neural data is available in the SSM package: (https://github.com/lindermanlab/ssm)

#### Estimation of time constants

We estimated the time constant of each mode of linear dynamical systems using eigenvalues *λ_a_* of the dynamics matrix of that system, derived by Maheswaranathan et al., 2019 as:

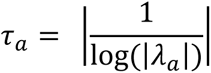

#### Decoding behavior from integration dimension

We trained a frame-wise decoder to discriminate pairs of behavior (such as sniffing vs attack) from the activity of the integration dimension on individual frames of a behavior (sampled at 15Hz) as described previously (Karigo et al., 2021). After merging bouts of behavior that were separated by less than 5 seconds, we trained a linear binary SVM, using 5-fold cross validation across intruder mice. Equal number of frames were used during decoder training to ensure chance decoder performance of 50%. ‘Shuffled’ decoder data was generated by training the decoder on the same neural data but with behavior annotations randomly assigned to each behavior bout. Decoding was repeated 20 times for each intruder and each mouse, and performance reported as the average accuracy across imaged mice.

#### Dynamic velocity as a measure of stability in a dynamical system and visualization as 3D landscape

We devised a metric termed the “dynamic velocity” to quantify the average intrinsically generated rate of change of the fit dynamical system during a given behavior of interest. We first calculated the average norm of *A_z_t__x_t_* for every value of *x_t_* associated with a given behavior, for a given state *z*. We then averaged this value across states, giving a definition of 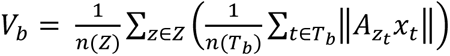, where Z is the set of states, *T_b_* is the set of all timepoints during which behavior *b* occurred, ‖·‖ is the Euclidean norm, and n(·) is the number of elements in a set. Finally, to facilitate comparison across animals, we normalized this value with respect to its maximum across behaviors in each animal.

We also converted the flow-fields obtained from rSLDS into a 3D landscape for visualization by calculating the dynamic velocity at each point in neural state space and using it as the height of a 3D landscape.

#### Statistical analysis

Data were processed and analyzed using Python, MATLAB, and GraphPad (GraphPad PRISM 9). All data were analyzed using two-tailed non-parametric tests. Mann-Whitney test were used for binary paired samples. Friedman test was used for non-binary paired samples. Kolmogorov-Smirnov test was used for non-paired samples. Multiple comparisons were corrected with Dunn’s multiple comparisons correction. Not significant (NS), P > 0.05; *P < 0.05; **P < 0.01; ***P < 0.001; ****P < 0.0001.

---

**Supplementary Figure 1:**
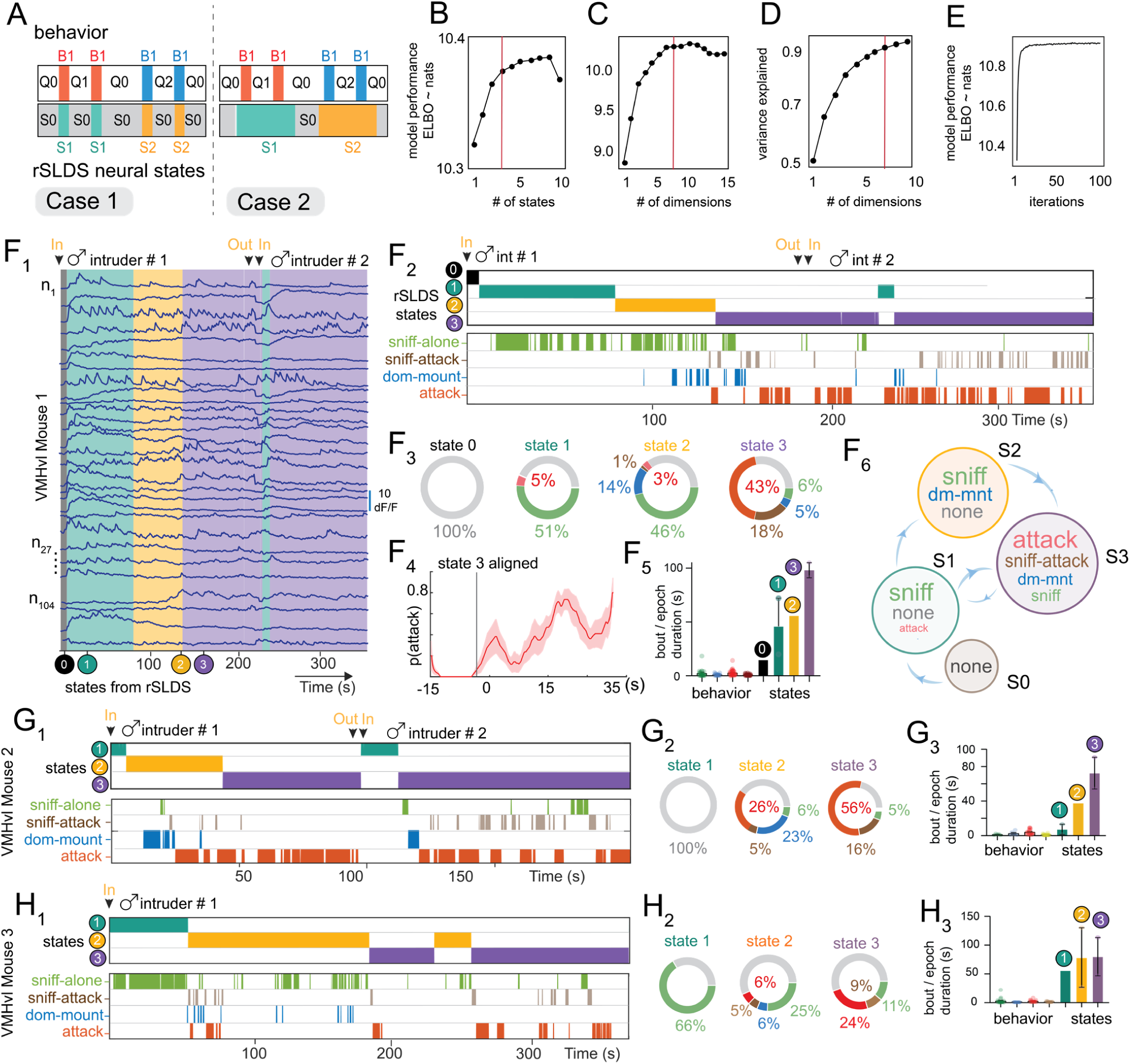
Unsupervised discovery of aggression-enriched states in VMHvl. Related to Figure 2 **A**: types of neural states that may be identified by rSLDS. B1, B2: behaviors; Q0, Q1: periods of quiescence between behavior bouts; S0,S1: rSLDS states discovered by analysis of neural data. Case 1 is when rSLDS states cannot distinguish behavior vs internal states. Case 2 is when rSLDS reflects internal state-encoding due to persistence during behavioral quiescence. **B**: optimization of number of states for rSLDS in example VMHvl mouse. Model performance is measured as the evidence of lower bound of the log-likelihood (ELBO). **C**: same as B, but for dimensionality. **D**: variance explained by dimension chosen in C. **E**: convergence of model performance. **F_1_**: rSLDS segments neural activity into various long-lived states in VMHvl mouse 1. **F_2_**: comparison of discovered states with behaviors performed by VMHvl mouse 1. **F_3_**: behavioral composition of discovered states highlights states with various amount of aggressive behavior. State 3 possesses the highest amount of attack behavior across mice (see panel I, N). **F_4_** : probability of attack aligned to the onset of state 3 across animals (n = 3 mice). **F_5_**: timescale of behavior bouts compared to the that of discovered states epochs. **F_6_**: state transition diagram from empirical transition probabilities. **G**: Same as F_2_, F_3_, F_5_ but for VMHvl mouse 2. **H**: Same as F_2_, F_3_, F_5_ but for VMHvl mouse 3.

**Supplementary Figure 2:**
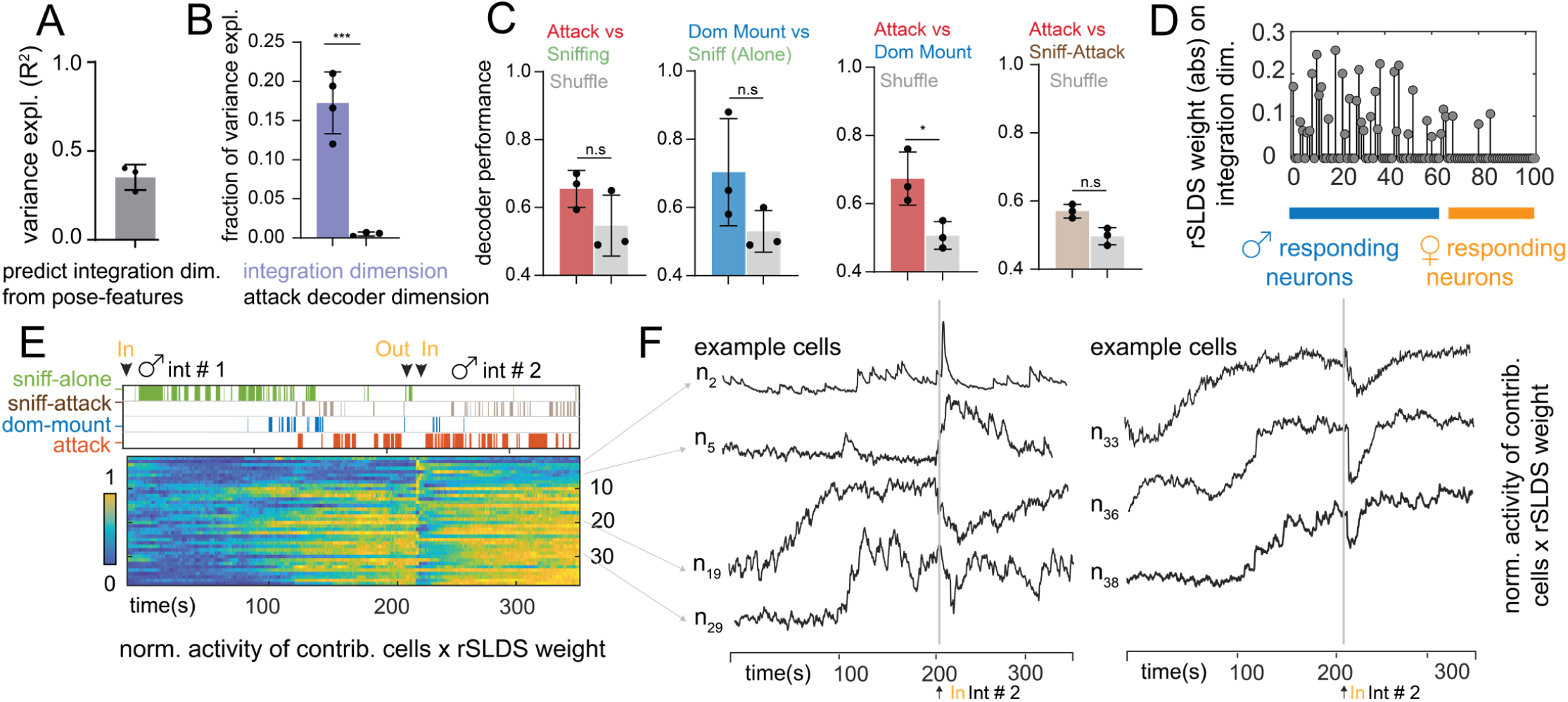
Characterization of aggression-integration dimension. Related to Figure 2 **A**: variance explained by GLM trained to predict integration dimension from pose-features including distance between mice, facing angle, speed, acceleration and velocity of resident mouse (mean: 0.35 ± 0.04 R^2^, n = 3 mice). **B**: fraction of overall variance explained by integration dimension (purple) compared to variance explained by decoder dimension trained to distinguish attack from sniff bouts (integration dimension mean: 0.17 ± 0.02, attack decoder mean: 0.005 ± 0.001, n= 4 mice, ***p<0.001). **C**: decoding behaviors from non-integration dimensions (average across dimensions, n = 3 mice). **D**: absolute rSLDS weight on integration dimension of VMHvl mouse 1, sorted by choice probability values for male vs female intruder encounter. **E**: normalized activity of neurons times rSLDS weight for cells with significant weights for integration dimension of VMHvl mouse 1. **F**: example cells from D.

**Supplementary Figure 3:**
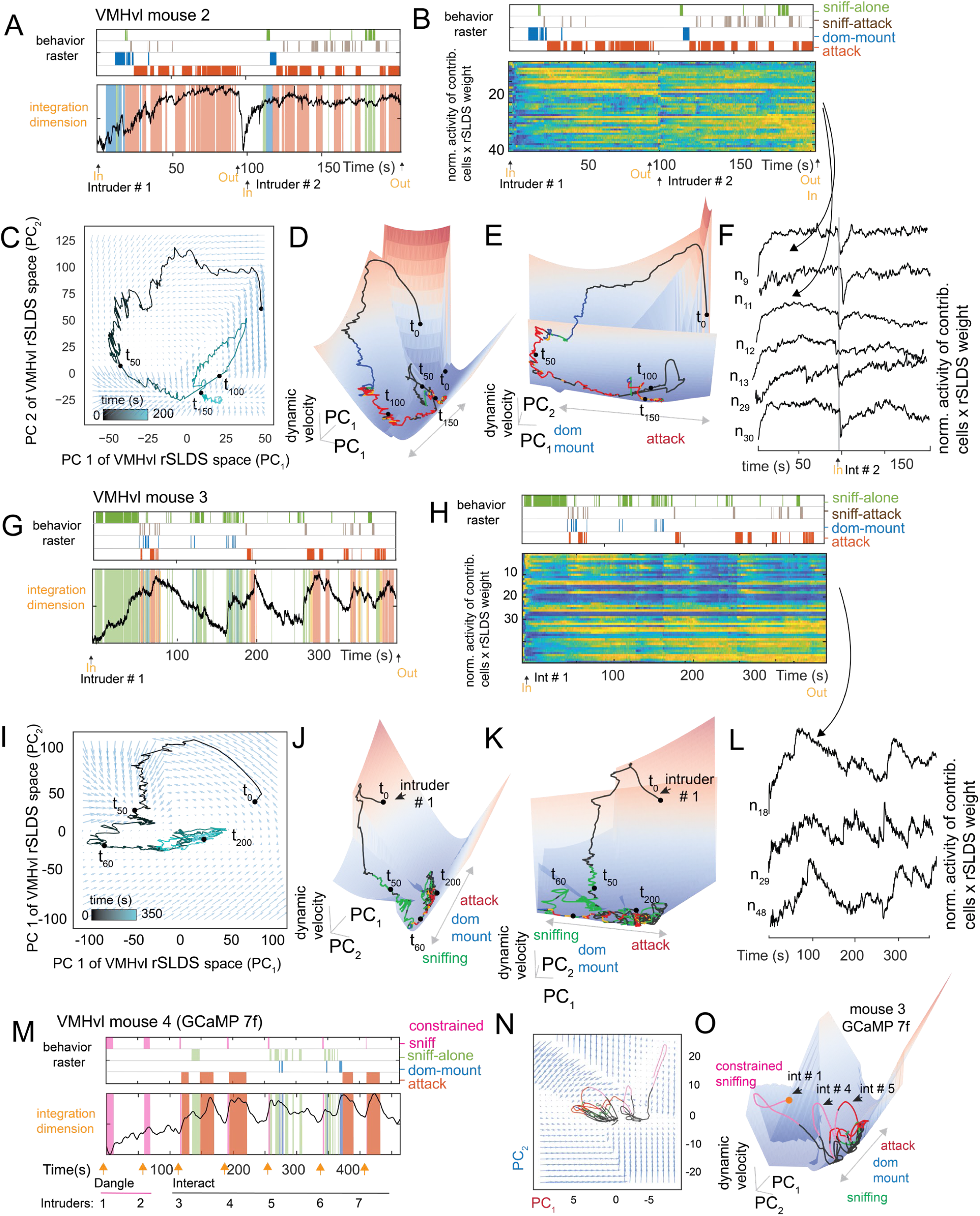
Discovery of approximate line attractor dynamics in VMHvl (mouse 2, 3 and 4) Related to Figure 3 **A**: integration dimension of VMHvl mouse 2. **B**: normalized activity times rSLDS weight for cells contributing significantly to integration dimension of VMHvl mouse 2. **C**: neural state space of VMHvl mouse 2 with population trajectory overlaid over inferred flow field. **D, E**: dynamic velocity landscape of VMHvl mouse 2 showing trough shaped landscape. **F**: example cell activity from B. **G-L**: Same as A-F but for VMHvl mouse 3. **M**: integration dimension of VMHvl mouse 4. **N**: neural state space of VMHvl mouse 2 with population trajectory overlaid over inferred flow field. **O**: dynamic velocity landscape of VMHvl mouse 4 showing trough shaped landscape.

**Supplementary Figure 4:**
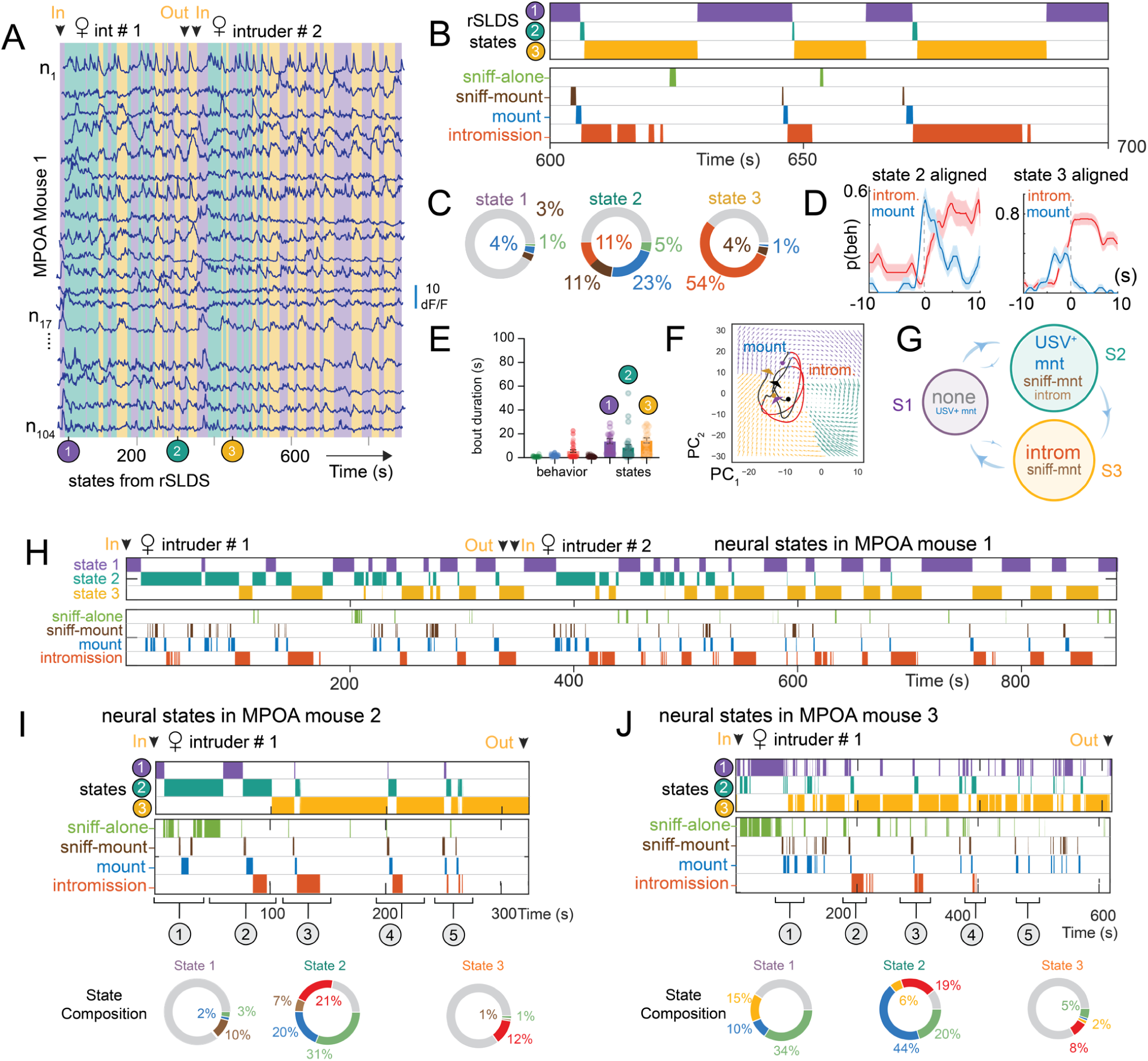
Unsupervised discovery of states by rSLDS in MPOA. Related to Figure 4 **A**: rSLDS segments neural activity into several short-lived states in MPOA mouse 1. **B**: states identified by rSLDS are closely aligned to behaviors performed by the resident mouse. **C**: behavioral composition of discovered states. **D**: probability of intromission and USV+ mounting aligned to the onset of state 2 and state 3 across animals (also see panel I, J, n = 3 mice). **E**: timescale of behavioral bouts and bouts of states. **F**: Same as Figure 4D but with state-specific inferred flow-field colors. **G**: state transition diagram from empirically calculated transition probabilities. **H**: state and behavior raster for MPOA mouse 1. **I**: state and behavior raster for MPOA mouse 2 (top). Behavioral composition of discovered states (bottom). **J**: same as I for MPOA mouse 3.

**Supplementary Figure 5:**
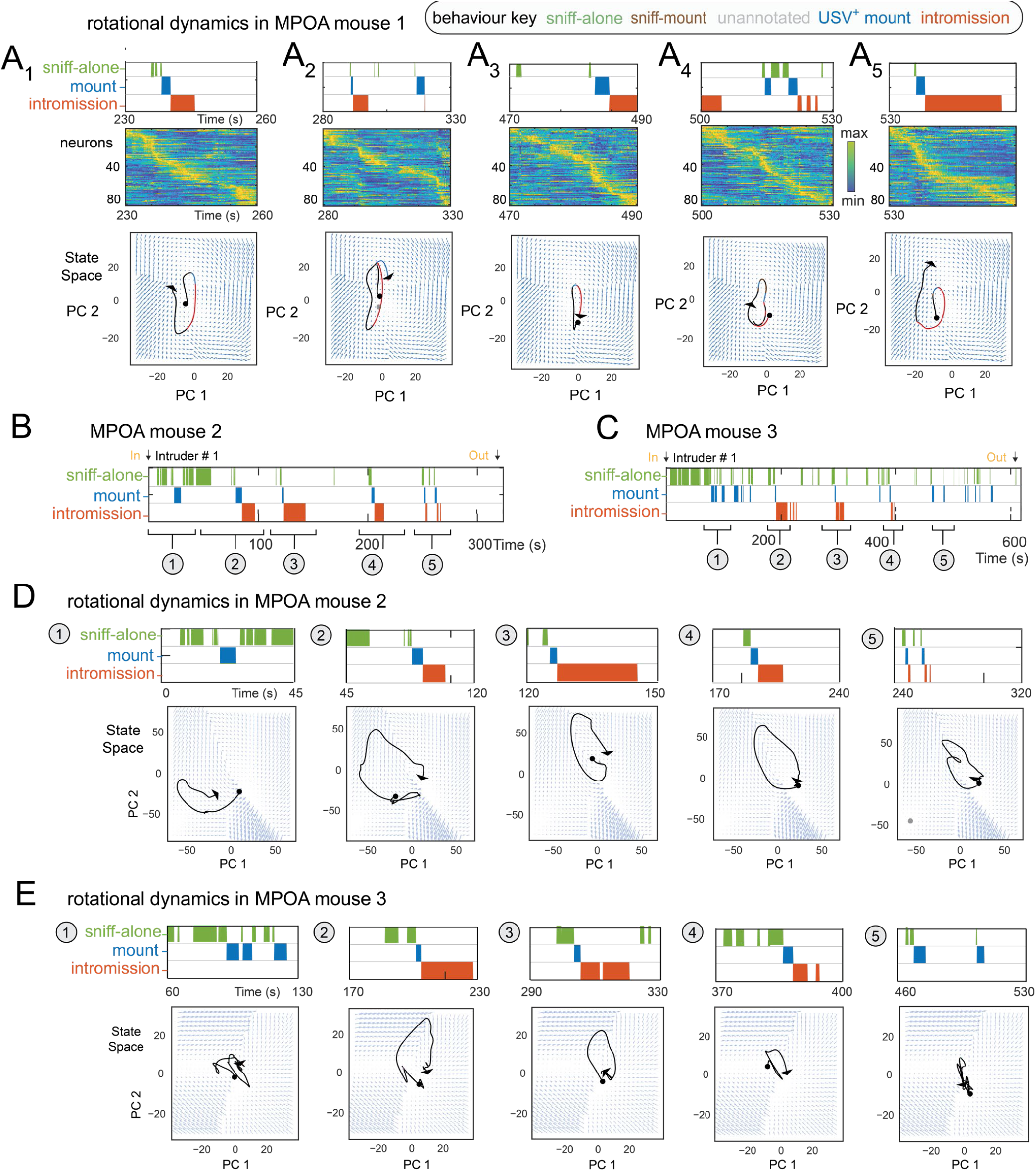
Rotational dynamics with sequential activity in MPOA. Related to Figure 4 **A_1_-A_5_**: Individual rotational trajectories for 5 mating episodes in MPOA mouse 1. **B,D:** Individual rotational trajectories for mating episodes in MPOA mouse 2. **C,E**: Rotational trajectories for mating episodes in MPOA mouse 3.

**Supplementary Figure 6:**
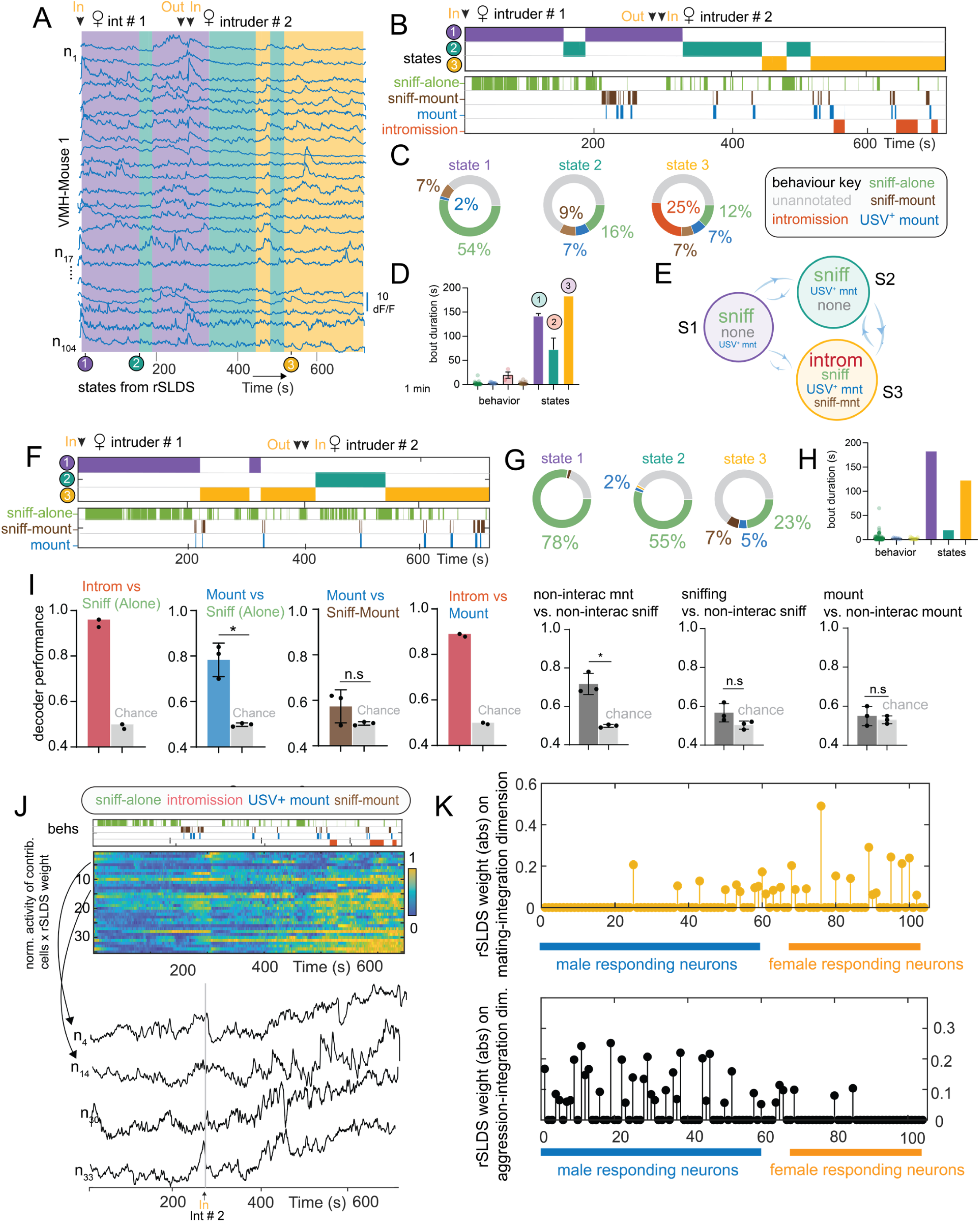
Unsupervised discover of mating-enriched states in VMHvl and characterization of mating-integration dimension. Related to Figure 6 **A:** rSLDS segments neural activity into various long-lived mating states in VMHvl mouse 1 during interactions with female intruders. **B**: comparison of discovered states with behaviors performed by VMHvl mouse 1. **C**: behavioral composition of discovered states highlights states with various amount of mating behavior. State 3 possesses the highest amount of mating behavior across mice (see panel H). **D**: timescale of behavior bouts compared to the that of discovered states epochs. **E**: state transition diagram from empirical transition probabilities. **F-H**: Same as B-D for VMHvl mouse 2. **I**: decoding behaviors from integration dimension (*p<0.01, n = 2 mice for intromission vs sniffing and intromission vs mounting, n = 3 mice for all other comparisons).). **J**: normalized activity times rSLDS weight for cells contributing significantly to integration dimension of VMHvl mouse 1. **K**: absolute rSLDS weight on integration dimension of VMHvl mouse 1 during mating behavior (top, yellow dots) and aggression (bottom, black dots) sorted by choice probability values for male vs female intruder encounter.

